# Single locus phosphoproteomics reveals phosphorylation of RPA-1 is required for generation of single-strand DNA following a break at a subtelomeric locus

**DOI:** 10.1101/2022.06.15.496243

**Authors:** Emilia McLaughlin, Annick Dujeancourt-Henry, Thibault Chaze, Quentin Giai Gianetto, Mariette Matondo, Michael D Urbaniak, Lucy Glover

## Abstract

Damage to the genetic material of the cell poses a universal threat to all forms of life. Central to the DNA damage response (DDR) is a phosphorylation signalling cascade that leads to the co-ordination of the cellular response to a DNA break. Identifying the proteins that are phosphorylated is crucial to understanding the mechanisms underlying this DDR. We have used SILAC-based quantitative phosphoproteomics to profile changes in phosphorylation site abundance following a single double strand break (DSB) at a chromosome internal locus and the subtelomeric bloodstream form expression site in *Trypanosoma brucei*. We report >6500 phosphorylation sites, including a core set of 211 DSB responsive phosphorylation sites. Along with phosphorylation of canonical DNA damage factors, we find that there is a striking distinction between the proteins phosphorylated in response to a chromosome internal DSB and one at the active BES and describe a single phosphorylation event on Replication factor A (RPA) 1 that is required for efficient resection at a bloodstream form expression site.

## Introduction

One of the most toxic insults to the genome is a DNA double strand break (DSB), where breaks occur simultaneously in the phosphate backbone of two complementary DNA strands. DSBs in the DNA can arise due to endogenous processes in the cell, such as replication fork collapse or stalling, and can also result from exogenous agents, such as chemicals or ionising radiation (Mehta and Haber 2014). DSB repair (DSBR) is a coordinated program of events initiated by a signalling cascade, which protein phosphorylation lies at the heart of. DSBs are detected by the MRN complex (MRE11-RAD50-NBS1) (Marechal and Zou 2013), and subsequently ATM (ataxia-telangiectasia mutated) and ATR kinase (ATM and Rad3-related), the master regulators of the DNA damage response (DDR), are recruited to the site of damage (Marechal and Zou 2013). This initiates a phosphorylation cascade in the cell with some 900 substrates modified (Matsuoka et al. 2007). One of the key substrates of ATM is the histone variant H2AX, where S139 is phosphorylated (termed γH2AX), and this is an early marker of the DSBR in mammals (Rogakou et al. 1998; Xiao et al. 2009). γH2AX spreads along large regions of chromatin fibre bi-directionally from the DSB site (Matsuoka et al. 2007) which aids the recruitment of chromatin remodelling factors, allows DNA damage proteins to access the DSB (Van and Santos 2018) and concentrates repair factors at the damaged site (Celeste et al. 2003).

Understanding phosphorylation cascades has been driven by Stable Isotopic Labelling of Amino acids in Cell culture (SILAC) (Ong et al. 2002) based quantitative phosphoproteomics (Bennetzen et al. 2010; Bensimon et al. 2010; Matsuoka et al. 2007; Zhou et al. 2016). SILAC phosphoproteomics has identified over 900 phosphorylation sites associated with ionizing radiation induced DNA damage, revealing a series of interconnected networks in the DDR including modules of proteins associated with DNA repair, replication, and chromatin modifications (Matsuoka et al. 2007). Over 70% of the phosphorylation sites identified are targets of the ATM kinase, and many of these substrates are themselves kinases, highlighting the central role of the phosphorylation cascade in the DDR.

Human African Trypanosomiasis (HAT) is a fatal vector borne disease caused by the protozoan parasite *Trypanosoma brucei.* In the mammalian host the parasite is found in the bloodstream, adipose tissue (Trindade et al. 2016) and skin (Caljon et al. 2016; Capewell et al. 2016). Here, the parasite is exposed to attack by the host immune system and is protected by a dense variant surface glycoprotein (VSG) coat, which is periodically exchanged by antigenic variation (Cross 1975; Horn 2014). *VSG’s* are exclusively expressed from one of 15 subtelomeric bloodstream form expression sites (BES) (Hertz-Fowler et al. 2008), however the majority of *VSG* genes are located in arrays in the subtelomeric regions of the megabase chromosomes, and also occasionally at chromosome internal regions (Berriman et al. 2005). There has been much debate into what triggers a *VSG* switch with DSBs (Boothroyd et al. 2009; Glover, Alsford, and Horn 2013), replication-derived fragility from the early replication of the BES (Benmerzouga et al. 2013; Devlin et al. 2016) or the formation of RNA:DNA hybrids (Briggs et al. 2018; Nanavaty et al. 2017) all being implicated. In *T. brucei* repair occurs predominantly via homologous recombination (HR) (Glover, McCulloch, and Horn 2008), but microhomology mediated end joining also makes a significant contribution at the active BES (Glover, Alsford, and Horn 2013). Within the HR pathway, sequence diversity amongst the genes facilitating repair suggesting functional divergence within the pathway as well (Dobson et al. 2011).

During the DNA damage repair cycle the G_2_/M checkpoint prevents division of unrepaired DNA and preserving genome integrity. Although trypanosomes do show cells arrested in G_2_/M following a DSB, some cells do continue to replicate and divide their DNA with a DNA break, which suggests a level of tolerance to DNA damage greater than that seen in other eukaryotes (Glover et al. 2007; Glover et al. 2019), perhaps aiding homology searching for antigenic variation. Early recognition and processing of a DSB via the MRN complex (<u>M</u>RE11, <u>R</u>AD50, <u>N</u>BS1 in mammals / XRS2 in *S. cerevisiae*) is important for both detection and signalling of a DSB. In *Trypanosoma brucei* and *Leishmania*, MRE11 maintains genomic integrity but does not affect the rate of VSG switching in the former (Laffitte et al. 2014; Laffitte et al. 2016; Robinson et al. 2002; Tan, Leal, and Cross 2002). However, RAD50 and MRE11 were shown to play distinct roles at the subtelomeric regions, with MRE11 required for efficient resection and both MRE11 and RAD50 promote recombination using longer stretches of homology (Mehnert et al. 2021).

Several DNA damage linked proteins directly influence antigenic variation. ATR mediates signal transduction in trypanosomes (Marin et al. 2020) and loss leads to increased VSG switching (Black et al. 2020), two RecQ-like helicases have been identified and mutants of one, RECQ2, show elevated *VSG* switching by telomere recombination and *VSG* gene conversion events (Devlin et al. 2016). RAD51, the primary recombinase in DNA repair, is required for homology searching and DNA strand exchange and is loaded onto single strand DNA by BRCA2, displacing RPA (Lo et al. 2003; Wong et al. 1997). In *T. brucei*, BRCA2 is essential for HR, DNA replication, cell division and antigenic variation (Hartley and McCulloch 2008; Trenaman et al. 2013), while RAD51 essential for HR and *rad51* null mutants have impaired, VSG switching (Conway et al. 2002; Glover and Horn 2014; Kim and Cross 2010; McCulloch and Barry 1999). In trypanosomes, five RAD51-related proteins DMC1, RAD51-3, 4, 5, and 6 are important for DSBR, but only RAD51-3 contributes to VSG switching (Proudfoot and McCulloch 2005). Antigenic variation occurs mainly by gene conversion (GC) events, where the active *VSG* is deleted and replaced by a silent donor (De Lange et al. 1983; Myler et al. 1985; Pays 1985; Robinson et al. 1999), but crossover switching events, where two *VSGs* are exchanged, have also been observed (Rudenko et al. 1996; Aitcheson et al. 2005; Pays 1985; Pays et al. 1983). The RTR complex which includes the RecQ-family helicase, a Topoisomerase IIIα, and RMI1/2 supresses these mitotic crossover and removes recombination intermediates (Mankouri and Hickson 2007). In trypanosomes *VSG* gene conversion and cross over events can be suppressed the RTR complex components *Tb*TOPO3α and *Tb*RMI1 or act in concert with RAD51 and RMI1 (Kim and Cross 2010, 2011).

Despite the importance of DSBR in evasion of the host immune system only one DNA damage associated phosphorylation site has been identified in *T. brucei*, that of γH2A (Glover and Horn 2012). In *T. brucei*, H2A Thr130 is phosphorylated in response to a DSB and colocalizes with RAD51 and RPA repair foci, typically during S or G2-phases of the cell cycle. The PTM of histones and non-histone proteins play a central role in the initiation and execution of the repair response (von Stechow and Olsen 2017; Price and D’Andrea 2013). In *T. brucei* histone acetyltransferase HAT3, which acetylates histone mark H4K4 (Siegel et al. 2008), and histone deactylase SIR2rp1 (Alsford et al. 2007) act in an acetylation and deacetylation cycle influencing RAD51 foci assembly and disassembly, respectively (Glover and Horn 2014). HAT3 also supresses VSG switching (Glover and Horn 2014) alluding to specific chromatin marks that regulate DNA recombination and VSG switching.

Here, use a quantitative single-locus phoshoproteomic approach to characterise changes in phosphorylation site abundance in response to a DSB at (i) a chromosome internal locus and (ii) the active BES. We found that there is a striking distinction between the proteins phosphorylated in response to a chromosome internal DSB and one at the active BES. Here we show that Histone H2A is phosphorylated at two additional positions in response to a DSB and that RPA-1 phosphorylation is differentially required for efficient resection at a chromosome internal versus subtelomeric break.

## Materials and Methods

### Trypanosome strains and culturing

*T. brucei* Lister 427 cell lines were grown in HMI-11 medium at 37.4 °C (74) with 5% CO_2_ and the density of cell cultures measured using a haemocytometer. Transformation of cell lines was carried out by centrifuging 2.5 x 10^7^ cells at 1000 g for 10 minutes at room temperature. The cell pellet was resuspended with 10 μg linearized DNA in 100 μl warm cytomix solution (75), placed in a cuvette (0.2 cm gap) and transformed using a Nucleofector^TM^ (Lonza) (X-001 function. Transfected cells were recovered in 36 ml of warm HMI-11 at 37 °C for 4 - 6 hours, after which cells were plated out in 48 well plates with the required drug selection. G418 selection was carried out at 2 μg/ml, puromycin at 2 μg/ml, blasticidin at 10 μg/ml and tetracycline at 1 μg/ml. Puromycin, phleomycin, hygromycin, blasticidin and G418 selection was maintained at 1 μg/ml. The VSG^up^ cell line has been described previously (Glover, Alsford, and Horn 2013) and the ^1^HR stain in (Glover, McCulloch, and Horn 2008).

### Generation of RPA-1 mutant cell lines

In order to generate the ^1^HRRPA-1^S5A^ cell line, the ^1^HR cell line was first transfected with 10 μg pRPA1^S5A^BLA, digested with NotI and KpnI and phenol/chloroform extracted. Successful integration of the construct was confirmed in the resulting clones by extraction of genomic DNA and amplification using the primer set RPA-1 5’ F primer and BlaNco1 R. The presence of the RPA-1^S5A^ mutation was confirmed by sequencing of the PCR product (Eurofins Genomics). Two correct single allele mutants’ clones were picked and then transfected with 10 μg pRPA1^S5A^NEO digested with NotI and KpnI and phenol /chloroform extracted. Genomic DNA was extracted from the resulting clones, and the mutant allele amplified using the primer set RPA1 5’F and Neo.2 R. The resulting PCR product was sequenced (Eurofins Genomics) to confirm the point mutation and a pair of biological clones used for downstream analysis. The VSG^up^RPA-1^S5A^ cell line was made in exactly the same way but using the VSG^up^ cell line.

### Quantitative single strand DNA resection assay

Quantification of ssDNA by qPCR was carried out using an adapted version of an established protocol (Mehnert et al. 2021; Zierhut and Diffley 2008). In brief, DNA was harvested at 0, 6 and 12 h following growth in 1 μg/ml tetracycline. 500 ng of extracted DNA was digested with either HindIII or mock digested (no enzyme) overnight at 37 °C. For the ^1^HR and ^1^HR RPA-1^S5A^ cell lines, qPCR was carried out using the RFP.2 F and RFP.2 R primers. For the VSG^up^ and VSG^up^RPA-1^S5A^cell lines, the qPCR reaction was carried out using the VSG21b F and VSG21bR primers. Luna Universal qPCR master mix (New England Biolabs) was used with 600 pM primer mix and 5 ng DNA per reaction. The PCR cycling conditions were: 95 °C for 3 minutes, and then 40 cycles of 95 °C for 10s and 55 °C for 30s on a thermal cycler. The Δc_q_ was calculated by subtracting the average c_q_ of the mock digest from the digested c_q_. % ssDNA was calculated using the formula: % resection = 100/[(1+2^Δcq^)/2], assuming 100% efficiency of the I-*Sce*I meganuclease. 3 technical replicates were carried out for each experiment and statistical analysis was carried out in Excel and GraphPad Prism version 9.

### VSG-seq analysis

For both the VSG^up^ and VSG^up^RPA-1^S5A^ cell lines, 5 x 10^7^ cells were harvested 0- and 7-days post growth in 1 μg/ml tetracycline, in triplicate, and RNA extracted. Frist strand synthesis was carried out using 500 ng of RNA, SuperScript® IV reverse transcriptase (ThermoFisher) and 200nM of the ‘All-*VSG* 3’-UTR’ primer (Mugnier, Cross, and Papavasiliou 2015) that binds specifically to the conserved 14-mer in the *VSG* 3’ UTR. The product was cleaned up using AmPureXP beads (Beckman Coulter). *VSG* cDNA was then amplified by PCR using 1 μg of cDNA, 0.2 mM dNTPs, 1 x PCR buffer, Phusion HF DNA polymerase (NEB), 200 nM of the spliced leader (SL) forward primer and 200 nM of the SP6-14mer reverse primer (Mugnier, Cross, and Papavasiliou 2015). The PCR conditions were: 5 cycles of: 94^0^c for 30s, 50^0^C for 30s and 72^0^C for 2 minutes, followed by 18 cycles of: 94 °C for 30s, 55 °C for 30s and 72 °C for 2 minutes carried out on a thermal cycler machine. The *VSG* PCR products were then cleaned up using AmpPureXP beads (Beckman Coulter) according to manufacturer’s protocols. Sequencing was carried out using a minimum of 4 μg product per sample at Beijing Genomics Institute (BGI) using the BGISEQ-500 platform in paired end mode. The number of million reads per library was: 4.01 for VSG^up^ uninduced replicate 1, 4.02 for VSG^up^ uninduced replicate 2, 4.00 for VSG^up^ uninduced replicate 3, 4.02 for VSG^up^ induced replicate 1, 4.00 for VSG^up^ induced replicate 2, 4.03 for VSG^up^ induced replicate 3, 3.90 for VSG^up^RPA-1^S5A^ uninduced replicate 1, 4.01 for VSG^up^RPA-1^S5A^ uninduced replicate 2, 4.00 for VSG^up^RPA-1^S5A^ uninduced replicate 3, and 4.01 for VSG^up^RPA-1^S5A^ induced replicate 1, 3.97 for VSG^up^RPA-1^S5A^ induced replicate 2 and 3.78 for VSG^up^RPA-1^S5A^ induced replicate 3. Reads were aligned to the *T. brucei* Lister 427 genome (Müller et al. 2018) with the BES’s and minichromosomal *VSGs* added from the VSGnome (Cross, Kim, and Wickstead 2014; Hertz-Fowler et al. 2008)) using Bowtie2 (Langmead and Salzberg 2012) with –very-sensitive parameters. BAM files were generated using samtools (Li et al. 2009). The overall alignment was: 98.82% for VSG^up^ uninduced replicate 1, 98.88% for VSG^up^ uninduced replicate 2, 99.13% for VSG^up^ uninduced replicate 3, 98.56% for VSG^up^ induced replicate 1, 98.32% for VSG^up^ induced replicate 2, 98.57% for VSG^up^ induced replicate 3, 99.06% for VSG^up^RPA-1^S5A^ uninduced replicate 1, 98.93% for VSG^up^RPA-1^S5A^ uninduced replicate 2, 98.89% for VSG^up^RPA-1^S5A^ uninduced replicate 3, and 98.19% for VSG^up^RPA-1^S5A^ induced replicate 1, 98.09% for VSG^up^RPA-1^S5A^ induced replicate 2 and 97.64% for VSG^up^RPA-1^S5A^ induced replicate 3. Reads aligning to each transcript were acquired using featureCounts (Liao, Smyth, and Shi 2014) and EdgeR (Robinson, McCarthy, and Smyth 2010) was used to perform differential expression analysis on all genes. The R script used to generate volcano and genome plots and perform differential genome analysis can be found in the thesis annex figure 5 and is adapted from (Mehnert et al. 2021)

### HMI-9 SILAC medium

For all proteomic analysis, HMI-9 SILAC medium (Urbaniak, Martin, and Ferguson 2013)-L-Arginine and – L-Lysine (Gibco, reference 074-91211A) was used. One 17.91g pot used to make up 1L of medium by the addition of 900 ml H_2_O, 2 g sodium bicarbonate (Sigma) and 14 μl of beta-mercaptoethanol and the mixture stirred for 1 h at room temperature. The pH was adjusted to 7.3, and the medium filtered using a 0.2 μM filter. The following components were added to the filtered medium: 10 ml of glutaMAX (ThermoFisher), 100 ml filter dialysed SILAC FBS (3 kda molecular weight cut off) (DC biosciences) Gibco, 5 ml Pen/ strep (5000 U/ml penicillin and 5000 μg/ ml streptomycin) and heavy or light labelled L-Arginine and L-Lysine to the concentrations stated in table 6 to make heavy and light medium, respectively.

**Table 1.** The concentration of light and heavy labelled L-Arigine and L-Lysine used to supplement both IMDM and HMI-9 SILAC medium.

| Amino acid | Label | Source | Mw | Final concentration in HMI-9 |
| --- | --- | --- | --- | --- |
| L-Arginine (R0) | Light | Sigma | 210.6 | 25.8 mg/l |
| L-Arginine (R6-<sup>13</sup>C<sub>6</sub>) | Heavy | Sigma | 216.6 | 25.9 mg/l |
| L-Lysine (K0) | Light | Sigma | 182.6 | 43.8 mg/l |
| L-Lysine (K4-<sup>2</sup>H<sub>4</sub>) | Heavy | Sigma | 223.1 | 53.5 mg/l |

### Assessing incorporation of the stable isotope label

To assess incorporation of the isotope label, the INT and VSG^up^ cell lines were seeded in ‘heavy’, and ‘light’ labelled SILAC HMI-9 and after 7 days 1 × 10^8^ cells harvested from each cell culture. Samples were extracted by FASP (see section 5.3) and processed for Mass Spectrometry as described below.

### Phosphoproteomic experimental set up

For preparation of samples for proteomic and phosphoproteomic analysis, SILAC adapted INT and VSG^up^ cell lines were seeded in SILAC ‘heavy’ and ‘light’ medium and cells grown in ‘heavy’ medium were induced using 1 μg /ml of tetracycline for 12 h. For each experimental condition, a label swap replicate was carried (induced cells grown in ‘light’ media and uninduced in ‘heavy’ media). Approximately 3 × 10^8^ cells were harvested from each culture condition by centrifugation at 1,000 × g for 10 minutes at 4 °C. The supernatant was removed, and the cell pellet resuspended in 200 μl ice cold PBS and transferred to a microcentrifuge tube, where it was centrifuged at 12,000 × g for 15s and the supernatant discarded. The cell pellet was lysed at 0.5 × 10^9^ cells/ml in ice-cold lysis buffer (0.1 mM TLCK, 1 μg/ml Leupeptin, 1x Phosphatase Inhibitor Cocktail II tablet (Calbiochem), 1 mM PMSF, 1 mM Benzamidine) and incubated at room temperature for 5 min, with cell lysis was verified by microscopy. Cell lysates were then stored at −80 °C before further processing.

### FASP protocol

The preparation of peptides for Mass Spectrometry (MS) analysis was carried out using FASP (Wisniewski et al. 2009) that has been optimized for *T. brucei* (Urbaniak et al. 2013). For the four samples generated for investigation of the DSB response, the total amount of protein concentrate was 1 mg and 2 mg of protein for each of the ^1^HR label swap replicates, and 0.89 and 0.62 mg of protein for each of the VSG^up^ replicates. For digestion of peptides, the concentrated sample was removed from the ultra-centrifugal filter and a 1:50 ratio of mass spectrometry grade Trypsin Gold (Promega) added to the sample which was incubated with shaking for > 12 hours at 37 °C in a thermal heat block. Trypsin digestion was inhibited by adding 0.1% formic acid (FA) to the digest.

### Peptide desalting

Peptides were desalted using a Sep-Pak C18 SPE cartridge (Waters) using a vacuum manifold according to manufacturer instructions. All buffers were freshly prepared. Briefly, C18 phase (Sep-Pak, Waters) was activated in methanol, rinsed once in 80% ACN 0.1% FA, washed thrice in 0.1% FA. The sample was then loaded onto the cartridge twice. Resin was washed thrice in 0.1% FA and peptides were eluted in 50% ACN 0.1% FA. The resulting sample was dried in a SpeedVac vacuum concentrator (ThermoFisher) until 50 µl remained and then transferred to a nano LC tube (ThermoFisher) and dried by lyophilization.

### Phosphopeptide enrichment

Phosphopeptide enrichment and Mass Spectromtery (MS) was carried out at the Mass Spectromtery for Biology Utechs (MSBio) platform at Institut Pasteur. Phosphopeptide enrichment was performed using a GELoader spin tip using EmporeTM C8 (3M) prepared for StageTip (Rappsilber, Mann, and Ishihama 2007) and washed sequentially with 100% MeOH and 30% ACN, 0.1% trifluoroacetic acid (TFA). Before the enrichment step, 10 mg/mL TiO_2_ slurry (Sachtopore-NP TiO_2_, 5 μm, 300 Å, Sachtleben) was prepared in 30% ACN, 0.1% TFA and introduced into the GELoader C8 spin tip. The spin column was packed by centrifugation at 100 × g and then equilibrated loading buffer (80% ACN, 6% TFA, 1 M glycolic acid) before loading lyophilised tryptic peptides resuspended loading buffer at a ratio of 1:5 peptides to beads. An aliquot of tryptic peptides was retained for proteome analysis. TiO_2_ spin tip was first washed with 80 % ACN, 6% TFA and then with 50% ACN, 0.1% TFA at 200 × g. Phosphopeptides were eluted from TiO_2_ beads by transfer to into a new microcentrifuge tube containing 20% FA, using 10 % NH_4_OH solution via centrifugation at 100 × g. To prevent the loss of phosphopeptides retained by the C8 plug a second elution was carried out with 80% ACN, 2% FA via centrifugation at 100 × g. Eluate fractions were combined and lyophilized prior to mass spectrometry analysis.

### Mass spectrometry analysis

Peptides were analyzed on a Q-Exactive HF instrument (Thermo Scientific) coupled with an EASY nLC 1200 chromatography system (Thermo Scientific). Samples were loaded at 900 bars on an in-house packed 50 cm nano-HPLC column (75 μm inner diameter) with C18 resin (3 μm particles, 100 Å pore size, Reprosil-Pur Basic C18-HD resin) and equilibrated in 98 % solvent A (H_2_O, 0.1 % formic acid) and 2 % solvent B (acetonitrile, 0.1 % formic acid). For both proteome and phosphoproteome analysis, peptides were eluted using a 3 to 29 % gradient of solvent B during 105 min, then a 29 to 56 % gradient of solvent B during 20 min and finally a 56 to 90 % gradient of solvent B during 5 min all at 250 nl/minute flow rate. The instrument method for the Q-Exactive HF was set up in the data dependent acquisition mode. After a survey scan in the Orbitrap (resolution 60,000), the 12 most intense precursor ions were selected for higher-energy collisional dissociation (HCD) fragmentation with a normalized collision energy set up to 26. Precursors were selected with a window of 2.0 Th. MS/MS spectra were recorded with a resolution of 15,000. Charge state screening was enabled, and precursors with unknown charge state or a charge state of 1 and >7 were excluded. Dynamic exclusion was enabled for 30 sec. For phosphoproteomic analysis, technical replicates were carried out in which samples acquisition was repeated twice for each label swap replicate.

### Mass spectrometry data processing

All raw data were searched using MaxQuant software version 1.6.1.0 (Cox and Mann, 2008; Tyanova et al., 2016), which incorporates the Andromeda search engine (Cox et al. 2011), against the *Trypanosoma brucei* 927 genome downloaded from TritrypDB (http://www.tritrypdb.org/) (Version 37, 11,074 protein sequences) (Amos et al. 2022) supplemented with frequently observed contaminants (such as mammalian keratins, porcine trypsin and bovine serum albumins). All SILAC features were selected by default using the appropriate heavy K and R amino acid to be detected. Modifications included carbamidomethylation (Cys, fixed), oxidation (Met, variable) and N-terminal acetylation (variable) and phosphorylation (S, T, Y variable). The mass tolerance was set to 6 parts per million (ppm) and peptides were required to be minimum 7 amino acids in length. Matching between runs allows peptides that are present in one sample but not identified by MS/MS in all samples to be identified by similarities in retention times and mass. The false discovery rates (FDRs) of 0.01 was calculated from the number of hits against a reversed sequence database. Only phosphorylation sites with a MaxQuant localisation probability > 0.95 were considered.

### Statistical analysis of proteomic data

Statistical analysis was carried out using Persues (Tyanova et al. 2016) version1.6.1.3. SILAC ratios were transformed to Log_2_ and intensities to Log_10_. Values were subject to further quality filtering such that ratios with >100% variation between label swap replicates were removed, and the localization probability of each phosphorylation site was required to be ≥ 0.95. Significantly changing phosphorylation sites were identified using Significance B testing (Cox and Mann 2008) which takes into account the intensity-weighted significance and used a Benjamini-Hochberg correction (Benjamini and Hochberg 1995) to set the false discovery rate at ≤ 0.01. Categorical enrichment was calculated using a Fischer’s exact test with a false discovery rate (FDR) ≤ 0.01. Gene Ontology (GO) term enrichment was carried out on the significantly enriched phopshosites for the ^1^HR and VSG^up^ data sets, using a Fischer’s exact test (FDR ≤ 0.05). All other statistical analysis was carried out in Microsoft Excel and GraphPad Prism, version 9.

## Results

### Adaptation of the ^1^HR and VSG^up^ cell lines to SILAC medium

In order to study the cellular response to locus specific DSBs, we used two established cell lines, ^1^HR and VSG^up^, that contain the tetracycline inducible yeast I-*Sce*I homing endonuclease which induces a DSB in approximately 95 % of all cells (Glover, McCulloch, and Horn 2008; Glover, Alsford, and Horn 2013). The ^1^HR cell line contains an 18 bp I-*Sce*I heterologous recognition sequence (*Sce*R; Figure 1A) at an intergenic chromosome internal polycistronic transcription unit on one homolog of chromosome 11 (Glover, McCulloch, and Horn 2008) (Figure 1A), and the VSG^up^ strain harbours the *Sce*I upstream of the actively expressed *VSG-2* on chromosome 6a (Figure 1B)(Boothroyd et al. 2009; Glover, Alsford, and Horn 2013). We then set out to characterise changes in the phosphoproteome in response to DSBs induced at (i) chromosome internal locus (^1^HR cell line) and (ii) within the active BES (VSG^up^ cell line).

**Figure 1.**
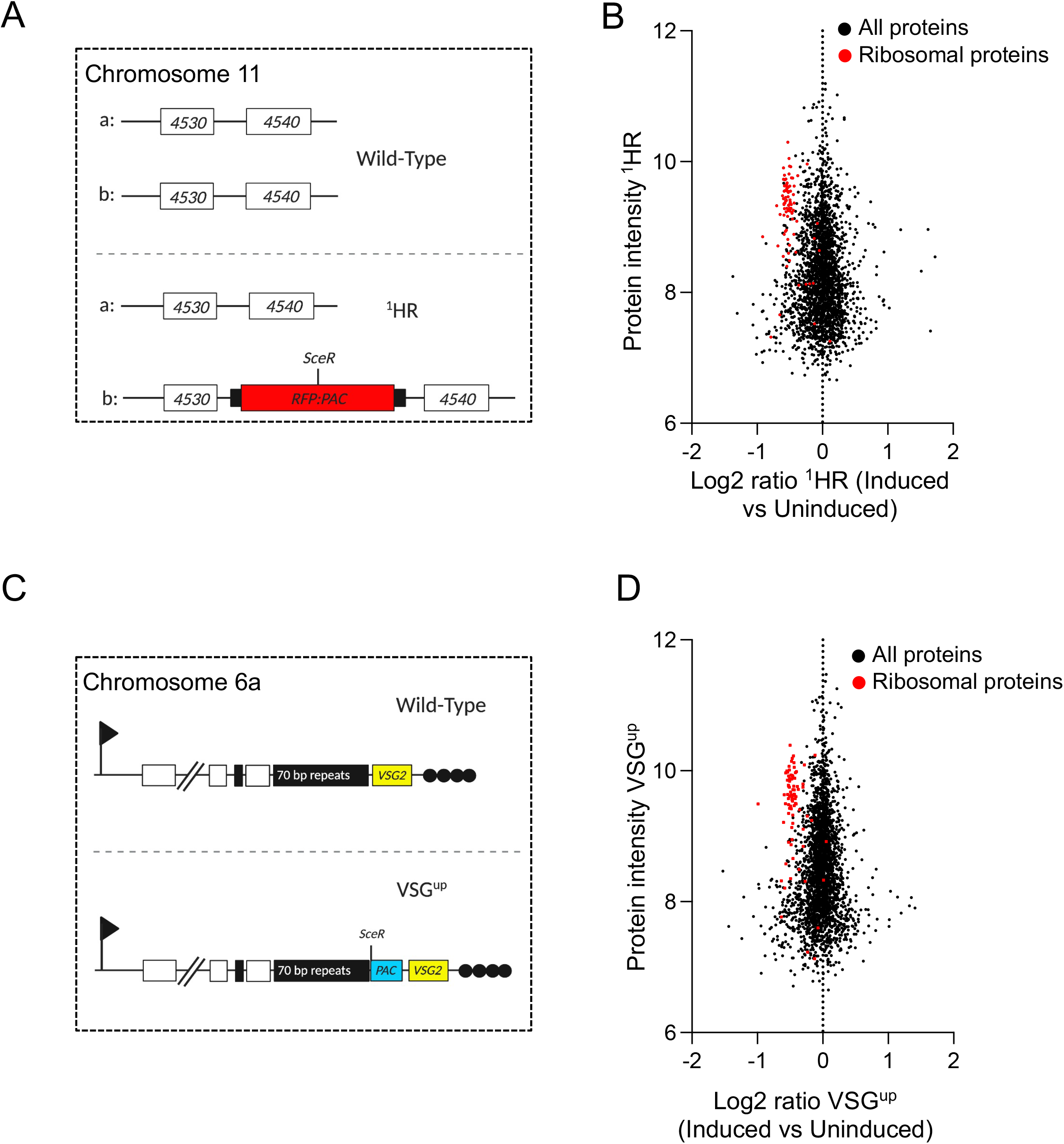
The ^1^HR and VSG^up^ DSB proteome. (A) Schematic of chromosome 11 Tb927.11.4530/40 locus, showing both alleles (upper panel: wild type) and then following modification to generate the ^1^HR chromosome-internal DSB cell line with the I-*Sce*I recognition site, SceR, highlighted (lower panel: ^1^HR). The DSB site is flanked upstream by *red fluorescent protein (RFP)* and *puromycin-N-acetyltransferase (PAC)* downstream. The site is positioned at an intergenic region between Tb927.11.4530 and Tb927.11.4540, shown as ‘4530’ and ‘4540’, respectively. Black boxes are tubulin intergenic sequences. (B) Total proteins identified in ^1^HR. The x - axis is the Log_2_ value of the ratio of each protein given as its presence in the DSB induced vs uninduced sample. The y-axis is the intensity of a given protein in the sample. Ribosomal proteins are highlighted in red, all other proteins shown in black. (C) VSG^up^ cell line set up showing the modified BES1 on chromosome 6a. An I-SceI meganuclease recognition site is inserted upstream of the actively expressed *VSG-2*, shown with a red vertical line. The I-SceI-R is flanked downstream by a *puromycin-N-acetyltransferase* gene (PAC). Arrow; native promoter of the expression site, white boxes; genes, solid black box; 70 bp repetitive sequence, black circles; telomere. (D) The VSG^up^ proteome, with details as described in (B). Black circle, individual proteins; red circles, ribosomal proteins. Schematics generated with BioRender.

SILAC experiments require the metabolic incorporation of stable isotope labelled amino acids present in the cell culture medium. Trypanosomatids are auxotrophic for arginine (R) and lysine (K) (Marchese et al. 2018) and we therefore used cell culture medium lacking in both and supplemented with either the ‘heavy’ isotope labelled L-Arginine U–^13^C_6_ and L-Lysine 4,4,5,5-^2^H_4_ (R_6_K_4_), or ‘light’ labelled L-Arginine and L-Lysine (R_0_K_0_). Trypanosome parasites have been shown to grow normally in SILAC HMI-9 and remain infective in mice (Urbaniak, Martin, and Ferguson 2013). Incorporation of labelled amino acids in the ^1^HR and VSG^up^ cell lines was assessed by mass spectrometry (MS), and we observed 93.4% and 96.1% heavy label incorporation in the ^1^HR and VSG^up^ cell lines, respectively (Figure S1A). In both the ^1^HR and VSG^up^ cell lines, γH2A foci form post DSB induction (Glover and Horn 2012; Glover, Alsford, and Horn 2013), we observed γH2A foci formation in cells grown in SILAC medium (Figure S1B), indicating a robust DNA damage response.

### Analysis of the total proteome following a double strand break

In bloodstream form trypanosomes, γH2A accumulation peaks at 12 h post DSB induction in both the ^1^HR and VSG^up^ cell lines (Glover and Horn 2012), and ssDNA accumulates between 9-12 h (Glover, McCulloch, and Horn 2008; Glover, Alsford, and Horn 2013). We therefore chose to carry out proteomic analysis at 12 h post DSB induction. For each sample, a label swap replicate was carried out (induced cells grown in ‘light’ media and uninduced in ‘heavy’ media). Analysis of the total peptide extract was carried out and in the ^1^HR data set, a total 2,457 proteins were identified (Figure 1C), and only 12 of these showed a > 2-fold change in abundance following DSB induction. In the VSG^up^ proteome, a total of 2,646 proteins were identified (Figure 1D) only 8 of which had a > 2-fold change in abundance following DSB induction. This confirms that large changes at the protein level are not seen at 12 h post DSB induction. However, we did observe a notable down-regulation of the ribosomal proteins in both the ^1^HR and VSG^up^ proteomes (Figures 1C and D). Ribosomal proteins are important for the assembly of ribosomal subunits and also function as RNA chaperones (X. Xu, Xiong, and Sun 2016). The specific down regulation of ribosomal genes observed here suggests that there a may be a global inhibition of protein translation in response to DNA damage as has been reported following a CRISPR-Cas9 induced DSB in mammalian cells (Riepe et al. 2021).

### The DSB locus-specific phosphoproteome of the ^1^HR and VSG^up^ strains

Within the total protein extract, phosphorylated peptides are of low abundance, and so we carried out an enrichment step using affinity purification on a TiO_2_ column (Rappsilber, Mann, and Ishihama 2007). In total, we identified 6,905 phosphorylation sites in the ^1^HR phosphoproteome (Figure 2A) and 6,540 for VSG^up^ (Figure 2B). Of these sites, 5,991 were common to both the ^1^HR and VSG^up^ data sets, and 914 and 549 sites were unique to the ^1^HR and VSG^up^ respectively (Figure 2C). Using γH2A as a positive control for the phosphoproteome analysis, in the ^1^HR strain we report an average of 5.39 fold increase in γH2A T131 (previously annotated as T130 excluding the initiator methionine (Glover and Horn 2012) (Figure 2A) and a 1.97 fold increase in the VSG^up^ strain (Figure 2B), confirming that we are able to detect DSB specific phosphorylation events using quantitative phosphoproteomics.

**Figure 2.**
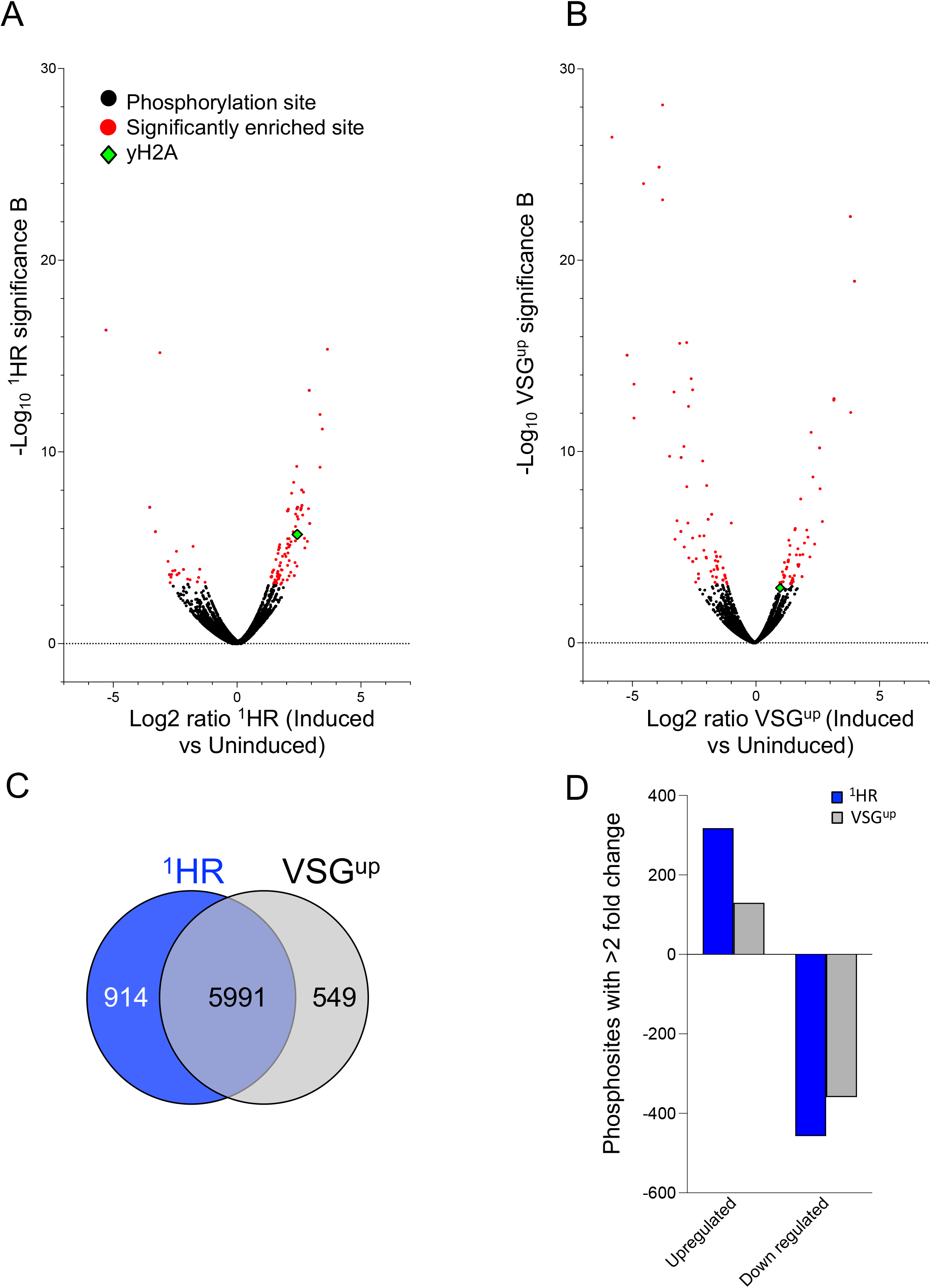
Locus specific phosphoproteome. (A) Quantification of changes in phosphorylation in the ^1^HR phosphoproteome. Black circles, non-significant change in phosphorylation; blue circles, significantly enriched phosphorylation sites; green triangle, γH2A. (B) Quantification of changes in phosphorylation in the VSG^up^ phosphoproteome, details as in (A). (C) Total number of phosphosites observed in ^1^HR and VSG^up^ phosphoproteomes, with those sites identified in both data sets shown. (D) Comparison of phosphorylation sites identified in the INT and VSG^up^ phosphoproteomes. Sites with over two-fold change in phosphoproteomic analysis are shown.

To identify significantly changing phosphorylation sites in the two data sets, we used Significance B testing (Cox and Mann 2008), which takes into account the intensity-weighted significance. Overall, there was no significant change in the distribution of phosphorylation events amongst phospho serine and thereonine, but our data set included no phosphorylation of tyrosine (Figure S2A) compared to published data. The majority of the phosphorylation events identified were on known proteins (Figure S2B). We identified 128 significantly altered phosphosites on 81 proteins in the ^1^HR phosphoproteome, 107 of which were upregulated and 22 downregulated (Figure 2A, Supplementary dataset 2). In the VSG^up^ phosphoproteome 135 significantly altered sites were identified on 95 proteins, 65 of which were upregulated and 70 down regulated (Figure 2B, Supplementary dataset 2). Within the phosphoproteomes, 26 sites were significantly enriched in both the ^1^HR and VSG^up^ strains. Of these 26 phosphorylation sites, 21 were upregulated in the ^1^HR, and only 3 were upregulated in VSG^up^, with down regulation of phosphorylation making a bigger contribution to subtelomeric DSBR. Amongst the 26 phosphosites that were significantly enriched in both the ^1^HR and VSG^up^ datasets, a number of modifications on proteins involved in RNA binding, RNA processing and translation were identified (Data set 1). We surveyed the 211 DSB responsive phosphorylation sites for categorical enrichment of Gene Ontology (GO) terms, using a Fischer’s exact test (FDR ≤ 0.05). The significantly enriched ^1^HR and VSG^up^ phosphorylation sites were further divided into phosphoproteins that were significantly upregulated or down regulated in each data set. In ^1^HR 6 significantly enriched GO terms were identified for upregulated phosphosites, which included histones and proteins that response to DNA damage (Figure S3A(i)), and one for VSG^up^ (Figure S3A(ii)). The reverse was seen with downregulated phosphosites; RNA binding was common to both, and 5 terms were enriched in the VSG^up^ including chromatin organisation (Figure S3B (ii)) and one for ^1^HR, RNA processing (Figure S3B (i)).

We next identified modifications on selected DNA repair proteins (Figure 3A). The H2A phosphorylation on a conserved S/T-Q motif (Redon et al. 2002; Rogakou et al. 1998) to give γH2AX, or γH2A in trypanosomes, is a widely studied early marker of DNA damage (Foster and Downs 2005; Glover and Horn 2012). We observe robust phosphorylation of *T. brucei* γH2A (T131) in response to a DSB (Figure 2A and B) and identified two additional modifications on the H2A C-terminus: S113 and S133 (Figure 3A), with only T131 and S133 being conserved amongst trypanosomatids (Figure 3B). Phosphorylation of S133 increased by 5.39-fold (p = 2.0283 x 10^-6^) in the ^1^HR strain and 1.97-fold (p = 1.3392 x 10^-3^) in the VSG^up^ strain (Data set 1). Individual phosphorylation events were detected on fragments harbouring exclusively T131 or S133, confirming that both sites are phosphorylated (Figure 3B). The third phosphorylation site identified on the H2A tail, S113, was 3.16-fold (p=0.000432) upregulated in response to an ^1^HR DSB and is conserved in *T. cruzi* but not Leishmania (Figure 3B) indicating that there are lineage specific modifications to the histone tail. In mammals, phosphorylation of the H2B C-terminus is a late marker of DNA damage, dependent on γH2AX (Fernandez-Capetillo, Allis, and Nussenzweig 2004). We detected a novel phosphorylation site at S39 on the C terminus of H2B (Tb927.10.10590) in ^1^HR strain (1.89-fold increase, p=0.0339), and in VSG^up^ (1.25-fold increase, p = 0.38) (Figure 3A). It is possible that H2B C-terminal phosphorylation will increase overtime until repair is complete. In mammalian cells 3 members of the NIMA (never in mitosis gene a)-related (NEK) kinase family, NEK1, NEK1, NEK10 and NEK11 are involved in the DDR and are implicated in check point control following DNA damage (Chen et al. 2011). We saw a significant enrichment of the phosphorylation of the serine/ threonine protein kinase NEK17 (Tb927.10.5950) in response to an ^1^HR DSB with two sites on NEK17, S197 and T195, increase by 6.14-fold (p = 6.13622 x 10^-8^) (Data set 1). NEK17 kinase is therefore a possible candidate for implementing phosphorylation marks that are specific to ^1^HR DSBR.

**Figure 3.**
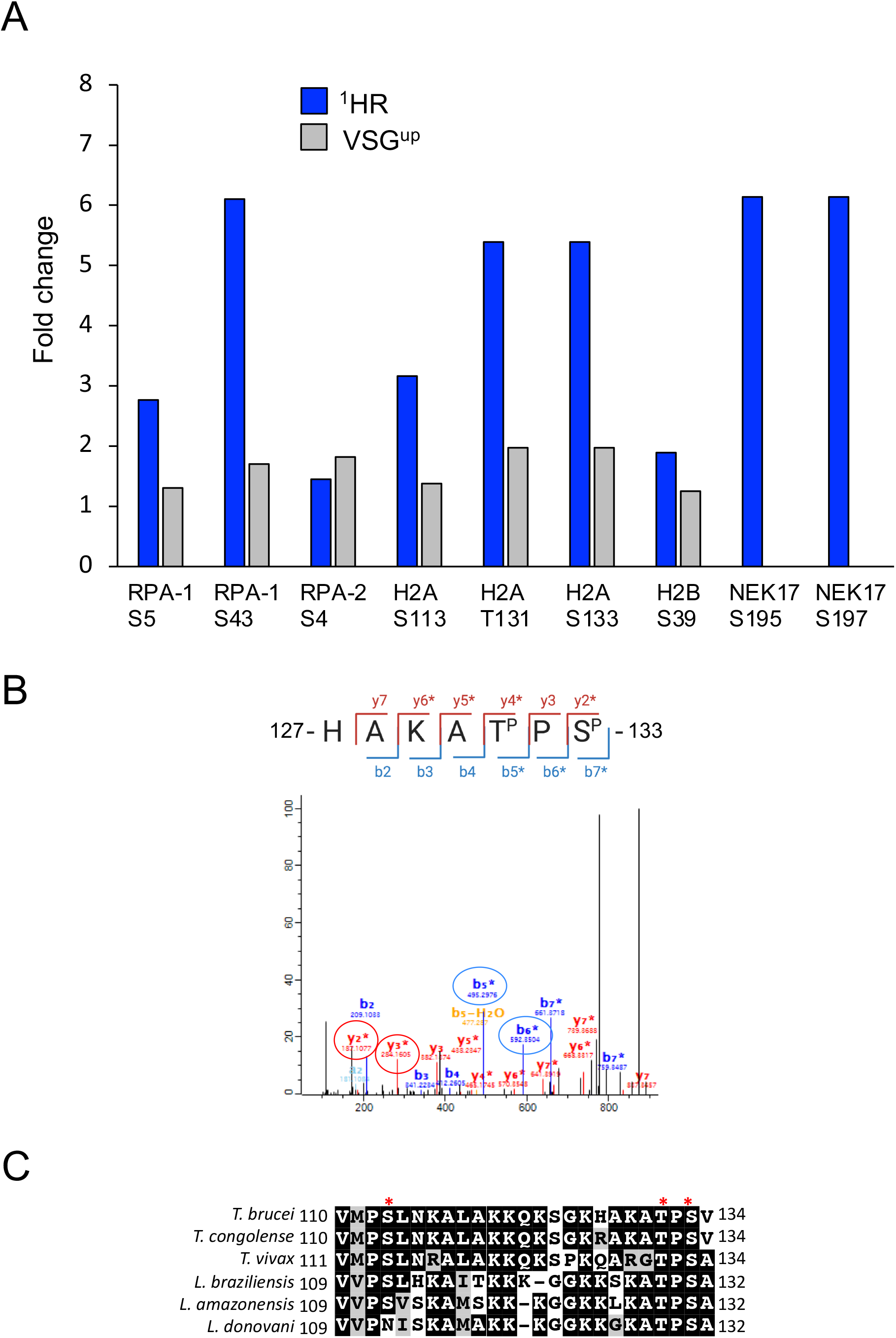
Phsophorylation of proteins involved DSBR. (A) Quantification of phosphorylation at specific amino acid sites. (B) Tandem mass spectral data showing the assignment of T131 and S133 phosphorylation sites on H2A (Tb927.7.2940). The MaxQuant localization score for each site 1. The loss of a phosphate group (−98 Da) is indicated with a *. The fragmentation pattern of the peptide is shown above. The b5 and b6 ions, highlighted on the spectrum with blue circles, show specific phosphorylation of T131, whilst the y2 and y3 ions highlighted with red circles. (C) Amino acid sequence alignment of the H2A C-terminus amongst trypanosomatids. Alignments shown are the c-terminus of H2A from *T. brucei* (Tb927.7.2940), *T. congolense* (TcIL3000_7_2140.1), *T. vivax* (TvY486_0702710), L. braziliensis (LbrM.32.1270, Leishmania amazonensis (LAMA_000111100) and Leishmania donovani (LdCL_210016600-t42). Red asterisks denote the S113, T131 and S133 phosphorylation sites; S, serine; T, threonine.

The heterotrimeric RPA complex, consisting of RPA-1, RPA-2 and RPA-3, is the major ssDNA binding protein in eukaryotes, and a number of phosphorylation sites of interest were identified on the complex in our data (Supplementary dataset 1). In mammals and yeast, the N-terminus of RPA-2 is hyper phosphorylated by ATR in response to a DSB (Maréchal and Zou 2015; Vassin et al. 2009). This hyperphosphorylation increases the affinity of RPA-2 for RAD51 (Wu et al. 2005) and promotes repair by HR (Shi et al. 2010). We identified only one phosphorylation site on RPA-2, S4, which showed a moderate up regulation of 1.45 and 1.82-fold in response to ^1^HR and VSG^up^ DSBs, respectively, suggesting that the N-terminus of RPA-2 is not hyperphosphorylated at 12 h post DSB induction (Figure 4A). Two sites on RPA-1 (Tb927.11.9130), the single stranded DNA binding component of the complex (Byrne and Oakley 2019), had specific sites phosphorylated. The first site, S5 was previously identified in a global phopshoproteomics analysis (Urbaniak, Martin, and Ferguson 2013) and we reported an average fold change in phosphorylation of 2.77 (p = 0.001) and 1.31 (p = 0.29) (Supplementary dataset 1) in response to DSBs induced in the ^1^HR and VSG^up^ cell lines, respectively (Figure 3A). The second site, S43 showed a 6.1-fold increase (p = 9.69 x10^9^) in response to an ^1^HR DSB (Figure 3A), and 1.7-fold increase (p = 0.06) in the VSG^up^ phosphoproteome (Supplementary dataset 1). Functional analysis of the *T. brucei* RPA-1 homolog (Tb927.11.9130) using InterPro (Blum et al. 2020) predicts 3 DNA binding OB fold domains (Figure S4A), similar to *L. amazonensis* RPA-1 (Pavani et al. 2014; Da Silveira et al. 2013) and *T. cruzi* RPA-1 (Pavani et al. 2016). The RPA-1 S43 modification lies in the first OB fold domain, whilst S5 lies outside of the OB fold domains (Figure S4A). In order to determine whether the phosphorylation sites identified are conserved amongst trypanosomatids and other eukaryotes we aligned the RPA-1 sequence of trypansomatids, *Homo sapiens* and *Saccharomyces cerevisiae* (Figure 1D). Multiple sequence alignment demonstrates that RPA-1 S5 lies in a trypanosomatid specific flexible linker, and the residue is conserved with *T. cruzi* but not *T. vivax* or *L. braziliensis*. The second phosphorylation site, RPA-1 S43, is conserved amongst *T. cruzi, T.* vivax and *L. braziliensis*, however not with *H. sapiens* or *S. cerevisiae* RPA-1 (Figure 4A). In *H. sapiens* the adjacent position, R42, has been identified as a key residue involved in interacting with DNA phosphates (Bochkarev et al. 1997).

**Figure 4.**
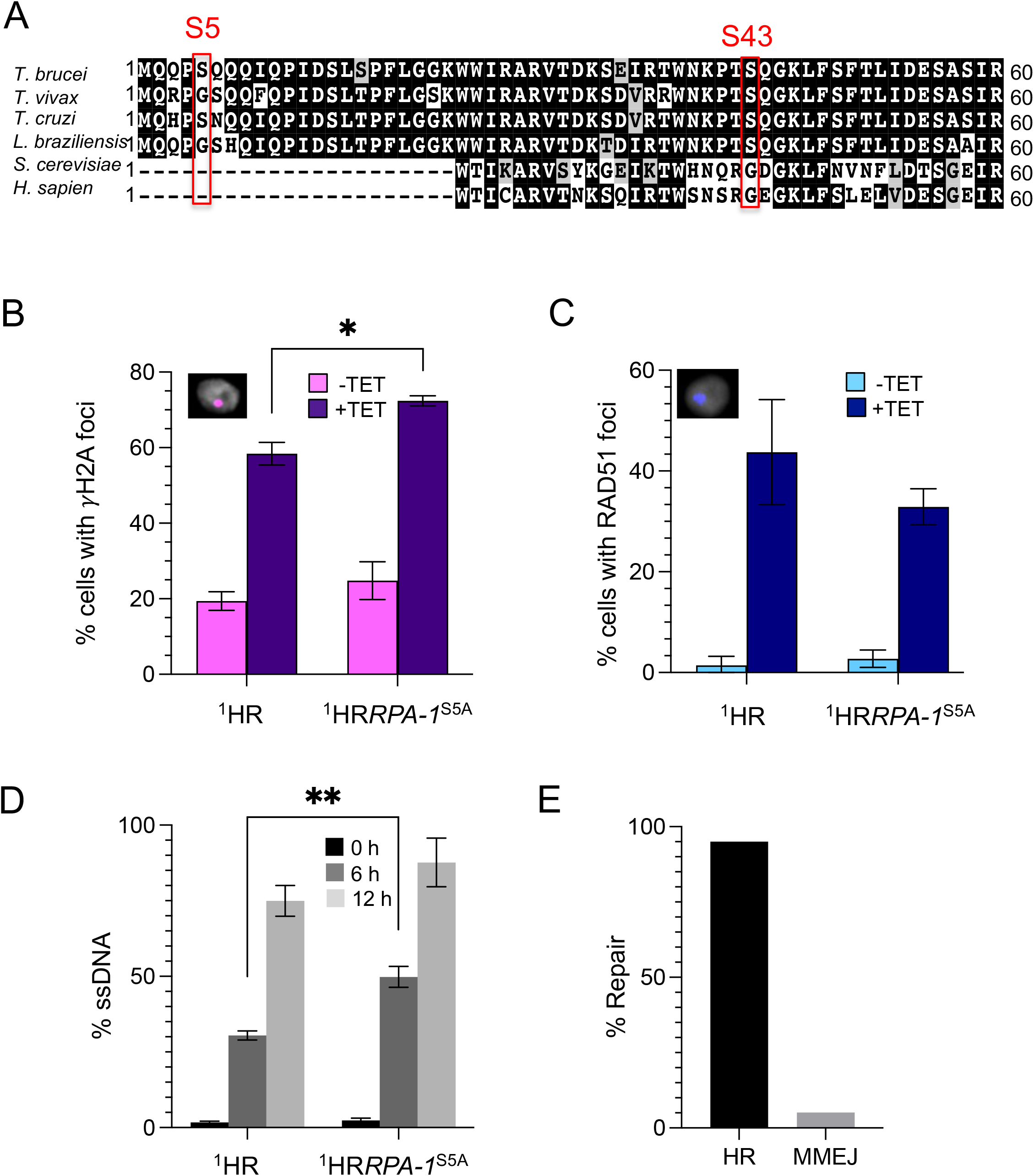
The RPA-1^S5A^ mutation increases yH2A DNA damage signalling and disrupts the timing of a chromosome internal DSB. (A) Amino acid sequence alignment of RPA-1 from *T. brucei* (Tb 927.11.9130), *T. cruzi* (TcCLB.510901.60) *T. vivax* (TvY486_1109660), *Leishmania brazillensis* (LbrM.28.1990), *Saccharomyces cerevisiae* (S288C) and *Homo sapiens* (GI: 4506583). Alignment to the first 60 residues of *T. brucei* RPA-1 is shown. Red boxes indicate phosphorylation sites identified in SILAC screens. (B) Monitoring of yH2A foci in the ^1^HR and ^1^HR*RPA-1*^S5A^ cell lines. n>100 for each sample. For the ^1^HR cell line error bars represent standard deviation between a pair of technical replicates. For the ^1^HR*RPA-1*^S5A^ cell line error bars represent standard deviation between a pair of biological clones. Significance was calculated using an unpaired t-test. Insert shows an example of a yH2A stained nuclei, with yH2A shown in magenta and DAPI in grey. (C) Monitoring of RAD51 foci formation in the ^1^HR and ^1^HR*RPA-1*^S5A^ cell lines, n> 100 per sample. Error bars for ^1^HR represent standard deviation of three technical replicates, and error bars for ^1^HR*RPA-1*^S5A^ cell line represent standard deviation between a pair of biological clones. Insert shows a representative RAD51 stained nuclei, with RAD51 foci shown in blue and DAPI in grey. (D) Percentage of ssDNA measured in the ^1^HR and ^1^HR*RPA-1*^S5A^ cell lines at 0, 6 and 12 h post DSB induction. For the ^1^HR cell line, the average and standard deviation of a pair of technical replicates is shown. For ^1^HR*RPA-1*^S5A^, the average and standard deviation of a pair of biological clones is shown. For all qPCR assays, each replicate is loaded three times on the qPCR plate. Significance is calculated using an unpaired t-test. (F) Percentage of repair events occurring by MMEJ or HR among 20 induced ^1^HR*RPA-1*^S5A^ subclones.

### The ^1^HR RPA-1^S5A^ mutant has increased γH2A signalling and a resection defect following a DSB

RPA-1 phosphorylation mediates ssDNA interactions in yeast (Yates et al. 2018), we therefore asked whether mutation of the phosphorylation sites identified in our phosphoproteomic data set led to a disruption of repair of a chromosome internal DSB (Glover, McCulloch, and Horn 2008). To determine the role of the phosphorylation of S43 we mutated this site to an alanine, which does not contain a hydroxyl functional group and therefore cannot be phosphorylated, in the ^1^HR cell line (Figure S4B). The replacement was confirmed by PCR in a pair of clones (Figure S4C and D). The resulting cell line is referred to as ^1^HR RPA-1^S5A^. Repeated attempts were made to mutate the S43 site, but no viable clones could be recovered even from a single allele replacement. This suggests that RPA-1 pS43 is essential for parasite growth and that disruption of the phosphorylation site is deleterious to parasite growth.

We observed a slight reduction in ^1^HR RPA-1^S5A^ cell growth over 96 h when compared to the parental ^1^HR strain, but growth was not significantly compromised (Figure S4E). In trypanosomes DNA damage can be monitored by both γH2A and RAD51 foci formation (Glover and Horn 2012). In the ^1^HR cell line γH2A foci were observed in 19.4% of uninduced cells had and this increased to 58.4% at 12 h post DSB induction, (Figure 4B) in agreement with previous findings (Glover and Horn 2012; 2014b; Mehnert et al. 2021). For the ^1^HR RPA-1^S5A^ mutants γH2A foci in uninduced cells were slightly increased at 24.5% and this rose to 72.4% following DSB induction, significantly higher than the induced ^1^HR strain (p = 0.0261) (Figure 4B) possibly reflecting unrepaired DNA lesions. We then looked at RAD51 foci formation in the ^1^HR cell line, foci were seen in 1.4% of uninduced cells, and this increased to 43.7% 12 h post DSB induction, consistent with previous reports (Glover, McCulloch, and Horn 2008; Mehnert et al. 2021). In the ^1^HR RPA-1^S5A^ cell line RAD51 foci were seen in 2.7% of cells and this increased to 32.9% at 12 h post DSB induction, less than seen for the ^1^HR cell line following a DSB (Figure 4C).

Repair of a DSB begins with 5’ to 3’ DNA resection either side of the break, catalysed by the MRE11 exonuclease, and the ssDNA overhangs serve as substrates for RAD51 filament formation and provides a template for HR (Syed and Tainer 2018). Given that γH2A signalling increased following a DSB in the ^1^HR RPA-1^S5A^ cell line, we asked whether the formation of ssDNA either side of the I-*Sce*I cleavage site is compromised by the RPA-1^S5A^ mutation. Using a quantitative resection assay (Mehnert et al. 2021) we found that ssDNA increased over 0 to 12 h post DSB induction with a relative percentage of 1.7%, 30.5, 75.0 % at 0 h, 6 h and 12 h, respectively (Figure 4D). The accumulation of ssDNA in ^1^HR cell line strain therefore follows a pattern similar to that observed previously using quantitative PCR (Mehnert et al. 2021) and slot blots (Glover, McCulloch, and Horn 2008; Glover and Horn 2014b). In the ^1^HR RPA-1^S5A^ cell line, the amount of ssDNA increased from 2.4% at 0 h to 49.8% at 6 h, and then to 87.7% at 12 h. The amount of ssDNA detected at all time points was increased in the mutant cell line, and this increase was significantly higher at 6 h post DSB induction (p = 0.002)(Figure 4D). This suggests that RPA-1^S5P^ restricts ssDNA resection at a chromosome internal site.

To determine if repair pathway choice was affected in the ^1^HR RPA-1^S5A^ cell line we generated a panel of subclones by clonogenic assays that were either uninduced (n = 5) or grown under DSB inducing condition (n = 11). The I-*Sce*I recognition site lies on an *RFP:PAC* fusion cassette (Figure 1A) in the ^1^HR cell line. Repair by homologous recombination will result in the loss of the fusion cassette, while repair by microhomology mediated end joining will result in retention of the fusion cassette but will result in a deletion mutation. Out of 11 induced subclones, only subclone was *RFP-PAC* positive, indicating that repair by HR predominates in the S5 mutant cell line (Figure 4D and Figure S5A). Sequencing of the region accommodating the *RFP-PAC* cassette was carried out for the one positive subclone, revealing an 81 bp deletion and therefore repair by MMEJ using short regions of homology.

### Mutation of S5 in RPA-1 leads to a resection defect following a subtelomeric DNA break

Our phosphoproteomic screens identified that RPA-1 S5 is phosphorylated in response to both chromosome internal (^1^HR) and telomeric (VSG^up^) DSBs (Figure 3A). To investigate repair at a subtelomeric region, we replaced both *RPA-1* alleles with a mutated *RPA-1^S5A^* as was described for the ^1^HR RPA-1^S5A^ cell line (Figure S6A and B). The replacement of both *RPA-1* alleles was confirmed by PCR (Figure S6A and B) and sequencing of the PCR product was used to confirm the S5A mutation. As in the ^1^HR RPA-1^S5A^ cell line, VSG^up^RPA-1^S5A^ showed only a slight reduction in cell growth in both uninduced and DSB inducing conditions over 96 h (Figure S6C), indicating that RPA-1 pS5 is not essential for the repair of a telomeric DSB. We next looked at the accumulation of γH2A. In the VSG^up^ cell line γH2A foci were observed in 19.9% of uninduced cells and this increased to 45.4% at 12 h post DSB induction (Figure 5A), consistent with previous findings (Glover, Alsford, and Horn 2013; Glover and Horn 2014b; Mehnert et al. 2021). For the VSG^up^RPA-1^S5A^ mutant, the number of foci rose from 25% to 46.6% at 12 h post DSB induction (Figure 5A), showing no significant difference from the parental cell line. This suggests that the RPA-1^S5A^ mutation has a larger impact on DSB repair at a chromosome internal site compared to at telomeric location. We did not assess RAD51 foci formation in the VSG^up^ mutants as RAD51 foci are not observed in response to telomeric DSBs (Glover, Alsford, and Horn 2013).

**Figure 5.**
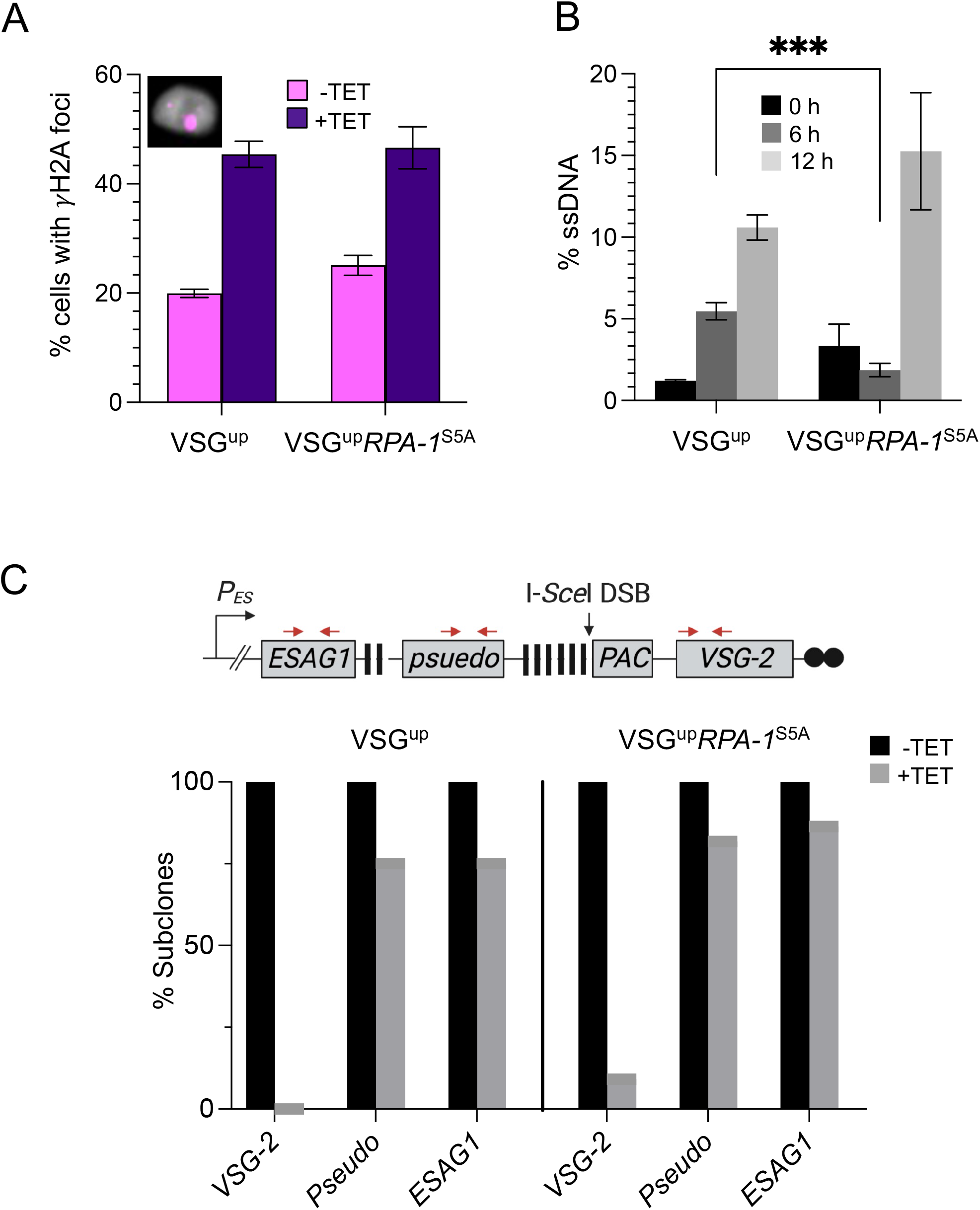
VSG^up^*RPA-1*^S5A^ mutants have a delay in ssDNA resection however repair at the active BES is not disrupted. (A) Monitoring of yH2A foci in the ^1^HR and ^1^HR*RPA-1*^S5A^ cell lines. n>100 for each sample. For the VSG^up^ cell line error bars represent standard deviation between a pair of technical replicates. For the VSG^up^*RPA-1*^S5A^ cell line error bars represent standard deviation between a pair of biological clones. (B) Percentage of ssDNA measured in VSG^up^ and VSG^up^*RPA-1*^S5A^ cell lines at 0, 6 and 12 h post DSB induction. For the VSG^up^ cell line, error bars represent the standard deviation of a pair of technical replicates and for INTRPA-1^S5A^ error bars represent the standard deviation between a pair of a pair of biological clones. Significance is calculated using an unpaired t-test. (C) PCR analysis of repair at the active BES of VSG^up^ and VSG^up^*RPA-1*^S5A^ sub clones. Upper panel, schematic of the modified BES1 with black arrows to show the position of primer binding sites used for subclone analysis. Lower panel, PCR analysis of VSG^up^ uninduced subclones (-Tet), n=5 and induced subclones (+ Tet) n=20. VSG^up^*RPA-1*^S5A^ uninduced sub clones n = 5, induced n=22

**Figure 6.**
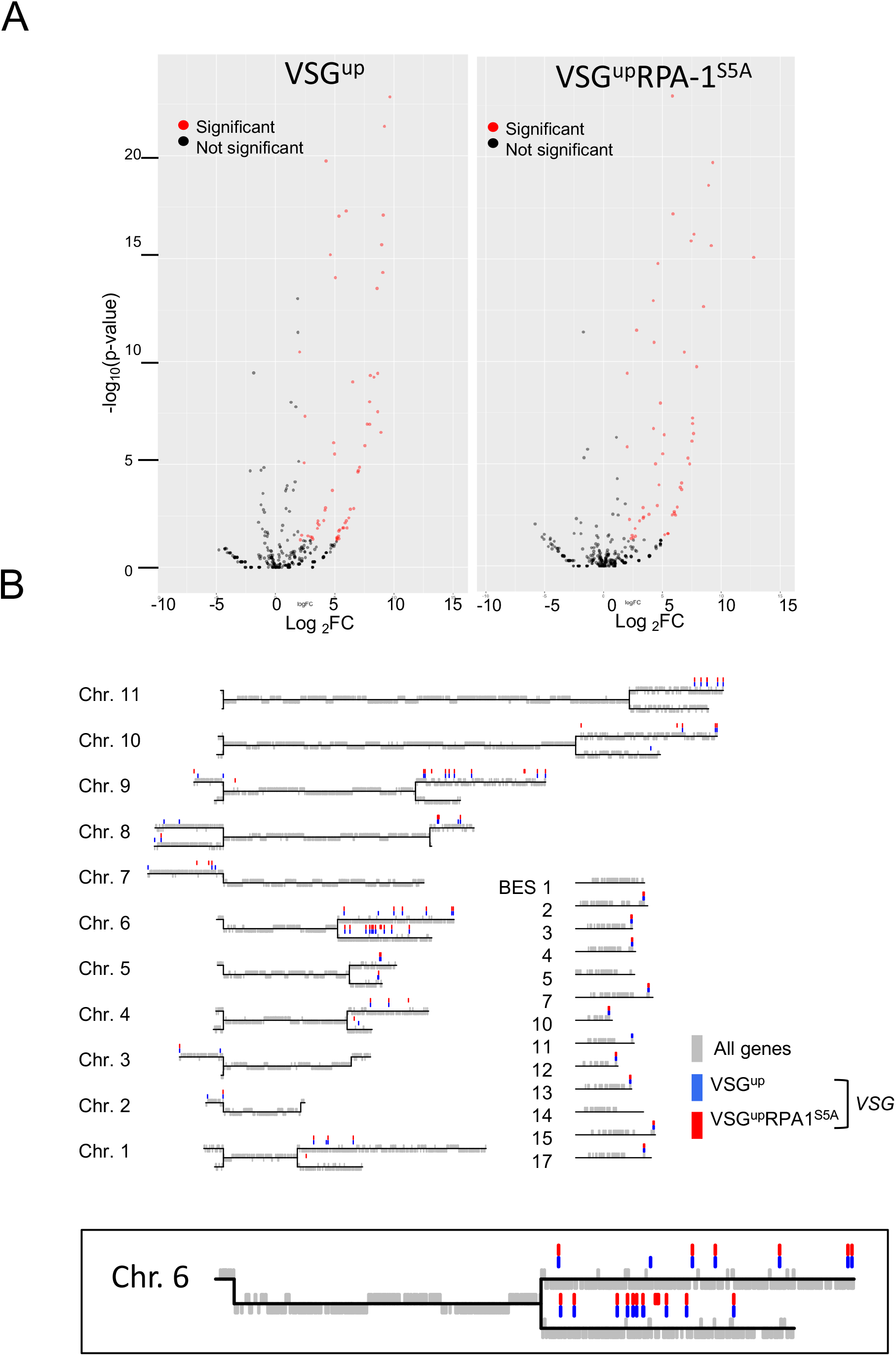
Antigenic variation is not compromised in the VSG^up^*RPA-1*^S5A^ mutants. (A) Volcano plots showing the significantly enriched *VSG* genes identified by VSG-seq following a BES DSB in the VSG^up^ and VSG^up^*RPA-1*^S5A^ cell lines. RNA was extracted 0 and 7 days following I-*Sce*I induction. Red circles represent genes that are significantly up regulated after 7 days induction (log_2_ FC >2 and P value < 0.05) and black circles represent genes that are not significantly up regulated. n = 3. (B) Map of the *T. brucei 427* genome showing the position of significantly up regulated *VSG* genes following a DSB. Megabase chromosomes are shown in black horizontal lines and genes in grey vertical lines. *VSG* genes that are significantly up regulated following a DSB are in highlighted with blue bars for the VSG^up^ and red bars for VSG^up^RPA-1^S5A^. Individual BES’s are shown separately on the right-hand side. An enlarged version of chromosome 6 is shown below.

We then assessed ssDNA accumulation in response to a subtelomeric DSB. Using our quantitative PCR assay (Mehnert et al. 2021) we determined that in the VSG^up^ cell line ssDNA accumulated over 12 h post DSB induction with a relative abundance of ssDNA of 1.2% at 0 h increasing to 5.5 % at 6 h and 10.6% at 12 h (Figure 5C) a pattern similar to that previously seen (Mehnert et al. 2021). In the VSG^up^RPA-1^S5A^ mutants, the relative amount of ssDNA was 3.3% at 0 h, and then decreased to 1.9% at 6 h, significantly lower than the same time point in the VSG^up^ cell line (p = 0.0007) and 15.3% at 12 h post DSB induction. The RPA-1^S5A^ mutation therefore disrupts ssDNA formation in response to a subtelomeric DSB with the initial stages of ssDNA resection inhibited by the mutation. This is in direct contrast to repair at a chromosome internal site, where the mutation leads to an increase in ssDNA resection. Given that the timing of ssDNA formation was disrupted in the VSG^up^RPA-1^S5A^ cell line, we were interested in whether VSG switching is disrupted following a DSB. Prior to DSB induction, 100% of the VSG^up^ population was VSG-2 positive by IFA, as expected (Figure S6D). Following a DSB in the VSG^up^ cell line, VSG-2 was detected in 6.7% of the population. The remaining 94.3% had replaced VSG-2, which we will refer to here as ‘VSG-X’. In the VSG^up^RPA-1^S5A^ mutants VSG-2 was also observed at the surface of all uninduced cells and this was reduced to 7.3% of the population upon DSB induction, with 92.3% of the population had switched to VSG-X (Figure S6D) suggesting VSG switching unaffected in the RPA-1^S5A^ mutants.

To investigate how repair comes about the in the RPA-1 mutants, we used a series of established repair assays (Glover, Alsford, and Horn 2013; Mehnert et al. 2021) to map individual repair events in a panel of induced and uninduced subclones generated for the VSG^up^ and VSG^up^RPA-1^S5A^ cell lines. BES1 contains 2 blocks of 70 bp repeats, the first block is located immediately upstream of the *VSG* and a second smaller 70 bp repeat region is found further upstream, flanked by *pseudo* gene and *ESAG1* (Figure 5C). The presence or absence of *VSG-2*, *ESAG1* and *pseudo* can be used to determine how repair has come about in induced subclones (Glover, Alsford, and Horn 2013). Repaired clones that retained *ESAG1* and *pseudo* but have lost *VSG-2* are expected to have undergone GC using the 70 bp repeats upstream of *VSG-2* as a homologous template. All uninduced subclones (n = 5 for both cell lines) were *VSG-2, ESAG1* and *pseudo* positive, as expected and *VSG-2* was lost in both VSG^up^ and VSG^up^RPA-1^S5A^ subclones (Figure 5C and S5A and B). All VSG^up^ induced subclones (n=20) and 20 out of 22 VSG^up^RPA-1^S5A^ had lost *VSG-2* (Figure 5C and S5A and B). The two *VSG-2* positive VSG^up^RPA-1^S5A^ clones were puromycin sensitive, indicating cleavage by I-*Sce*I, and have likely undergone repair by MMEJ. *ESAG1* and *pseudo* were retained in 75% of induced VSG^up^ subclones (Figure 5C), consistent with previous findings that GC using the large block of 70 bp repeats predominates in this cell line (Glover, Alsford, and Horn 2013), and 82% of VSG^up^RPA-1^S5A^ subclones (Figure S7B). Overall, we did not observe a considerable difference between repair at the BES in the parental and mutant cell line.

### The RPA-1^S5A^ mutation does not disrupt access to the *VSG* repertoire

The *T. brucei* genome contains over 2500 *VSG* genes and *pseudo* genes in subtelomeric arrays (Berriman et al. 2005; Cross, Kim, and Wickstead 2014), which can be used as templates for recombination during antigenic variation. The order in which these archival *VSGs* are activated is thought to be a semi-ordered process during an infection, with BES associated *VSG*s being most frequently activated followed by *VSGs* found in the subtelomeric arrays and finally *pseudo* genes (Morrison et al. 2005; Marcello and Barry 2007; Stockdale et al. 2008). We therefore assessed whether the RPA-1^S5A^ mutation disrupted access to the antigen repertoire during a *VSG* switch. To do this we used VSG-seq (Mugnier, Cross, and Papavasiliou 2015), which makes use of conserved sequences in the 5’ SL and 3’ UTR of every *VSG* mRNA to amplify and subsequently sequence of all of the *VSG*s expressed in a population. The expressed *VSGs* in the VSG^up^ and VSG^up^RPA-1^S5A^ cell lines were amplified and sequenced 0- and 7-days post DSB induction. *VSG* genes were considered significantly enriched when the log_2_ fold change (FC) of the induced (+ Tet) sample compared to the uninduced (-Tet) is > 2, with a p <0.05. We identified 136 *VSG* sequences that were significantly enriched in the VSG^up^ cell line (Figure 10A) and 128 were enriched in the VSG^up^RPA-1^S5A^ cell line (Figure 5D). We next looked at the genomic locations of the significantly enriched genes by mapping them to the *T. brucei* 427 genome (Müller et al. 2018) (Figure 5E). In the VSG^up^ cell line 10 BES *VSGs* and 27 *VSGs* located on the minichromosomes were enriched. In the VSG^up^RPA-1^S5A^ cell line, 9 BES *VSGs* were enriched and 25 minichromosomal *VSGs* (Figure 5E). Therefore, the RPA-1^S5A^ mutant does not significantly alter the number and genomic positions of the templates used to repair a BES DSB, with both cell lines using a similar number and variety of templates for DSBR

## Discussion

Here we report the use of SILAC quantitative phosphoproteomics to characterise the *T. brucei* DSB phosphoproteome in response DSBs targeted at both chromosome internal and subtelomeric loci. While the purpose of this study was to determine the specific phosphorylation events that govern the DDR in *T. brucei*, analysis of the proteome revealed that ribosomal proteins are down regulated following a DNA break. A similar phenomenon is seen in human cells where phosphorylation of eIF2a halts translation following a DSBs via ribosome remodelling (Riepe et al. 2021), whilst mouse embryonic fibroblasts exposed to global DNA damage results in transcriptional silencing in the nucleolus (Kruhlak et al. 2007). The down regulation of ribosomal proteins identified here alludes to cross talk between the nucleolus and the DDR, as has been observed in other organisms (Ogawa and Baserga 2017).

Our DNA damage phosphoproteome implicated RNA binding proteins (RBP) in DSBR. Phosphoproteomic studies of the DNA damage response in human cells have also identified enrichment of proteins with RNA binding capacity (Matsuoka et al. 2007; Bennetzen et al. 2010) and a number of RBPs have been shown to have a dual role in both RNA binding and the DDR. Such is the emerging evidence for the roles of RBPs in the DDR that a new class of proteins has been defined, the DNA damage response RNA binding proteins (DDRBPs) (Dutertre and Vagner 2017). Some of these DDRBPs can also bind double or ssDNA and are involved in regulating R-loop formation (Aguilera and Garcia-Muse 2012; Dutertre and Vagner 2017) by coating the nascent RNA (Nishida et al. 2017; Sollier et al. 2014). R-loop formation is associated with increased DNA damage and VSG switching in *T. brucei* (Briggs et al. 2018) and it is possible that some of the RBPs identified here supress R-loop formation following DSB induction. However, it is of note that in yeast R-loop formation is an important part of efficient HR (Keskin et al. 2014; Ohle et al. 2016), and it is also possible that RBPs assist in productive R-loop formation that contributes to DSBR.

The phosphorylation status of a given protein is a dynamic equilibrium balanced by the actions of protein kinases and protein phosphatases. In human cells, phosphoproteomic analysis of the DSBR revealed that approximately one third of the total sites identified are dephosphorylated in response to a DSB (Bennetzen et al. 2010; Bensimon et al. 2010), and here dephosphorylation was highly represented in VSG^up^, accounting for 51% of significantly altered modifications, indicating its important role at the subtelomeric locus. In contrast, the majority of significantly altered phosphorylation sites following an ^1^HR are upregulated, again highlighting the disparity between chromosome internal and subtelomeric repair. Histone modifications play a key role in the DNA damage response (DDR), regulating access to chromatin and signalling for DNA damage (Van and Santos 2018). Key to the DDR is the phosphorylation of histone H2A at T131 to give γH2AX in trypanosomes. In mammals, the γH2AX modification occurs at S139, however T136 and other phosphorylation sites on the histone tail have also been shown to be involved in the mammalian response to DNA damage (Redon et al. 2002; Xie et al. 2010). In *S. cerevisiae*, systematic mutation of the histone tail identified three sites on the H2A tail that are important for DNA damage, all of which are involved in DSBR by HR (Moore et al. 2007). In our phosphoproteome we identified two additional sites in histone H2A that are phosphorylated in response to a DNA break: S113 and S133. Given that these modifications were not previously annotated in global analysis of the *T. brucei* phosphoproteome (Benz and Urbaniak 2019; Urbaniak, Martin, and Ferguson 2013) they are likely highly specific to DSBR. We suggest that as the T131 lies four residues from the C terminus of H2A, it is the analogous modification to that seen in both mammalian and yeast DDR, however whether there is interdependency between the two sites requires further investigation.

One of the greatest up regulated phosphorylation sites following a ^1^HR DSB was the modification of RPA-1 S43, which increased by an average of 6.1-fold, suggesting that the protein is abundantly and specifically phosphorylated in response to a break. The RPA-1 S43 phosphorylation site identified here was not previously identified in a global study of the *T. brucei* phosphoproteome (Urbaniak, Martin, and Ferguson 2013) indicating that the modification is DNA damage specific. We were unable to generate homozygous RPA-1^S43A^ mutants despite repeated attempts and we speculate that that prohibiting the phosphorylation of RPA-1 S43 is lethal to the cell. RPA-1 S43 lies within the conserved DNA binding domain OB1, which has the highest affinity for ssDNA in trypanosomes (Neto et al. 2007; Pavani et al. 2016; Pavani et al. 2014) and is conserved amongst trypanosome species. The adjacent residue in *H. sapiens* RPA-1 has been shown to directly interact with DNA phosphates (Bochkarev et al. 1997) and it is therefore possible that disruption of the neighbouring residue in trypanosomatid RPA-1 destabilises the interaction with ssDNA. Given the diverse functions of the RPA complex in the cell (Byrne and Oakley 2019), the introduction of a mutation that disrupts ssDNA binding would likely be highly damaging to the cell.

The second phosphorylation site identified on RPA-1, S5, showed a moderate increase in phosphorylation compared to that of S43. RPA-1 S5 mutants showed an increase in γH2A foci formation and ssDNA at a chromosomal internal break, suggesting prolonged unrepaired damage, interestingly this increase was not reciprocated following a break at the active expression site. A greater increase in phosphorylation was identified following an ^1^HR DSB compared to a VSG^up^ DSB (2.7 FC in phosphorylation and 1.7 FC, respectively) which may be related to the strong preference for HR at this site (Glover, McCulloch, and Horn 2008), which involves the generation of long stretches of ssDNA either side of the DSB that are stabilised by the binding of the RPA complex. The RPA-1 S5 mutation had contrasting effects on ssDNA accumulation at a subtelomeric locus, here ssDNA accumulation severely reduced in the first 6 h following a break, indicating that the initial stages of resection are inhibited. DNA resection at the BES has previously been shown to be associated with the histone acetyl transferase HAT3 and TbMRE11, which supresses subtelomeric ssDNA formation (Glover and Horn 2014; Mehnert et al. 2021). Interestingly, despite the disruption in ssDNA accumulation at the subtelomeric loci, repair at the BES continues unrestricted, and cells were able to undergo recombination driven *VSG* switching. We did not see a change in the access to the genomic repertoire following a *VSG* switch, as has been reported upon *TbMRE11* and *TbRAD50* knockout (Mehnert et al. 2021). Unlike in mammalian cell, we did not detect hyperphosphorylation of RPA-2 N-terminal tail, rather a single phosphorylation event at S4. RPA-2 N-terminal phosphorylation is associated with checkpoint activation in mammals (Byrne and Oakley 2019), and it is possible that this signalling pathways is absent in trypanosomes. Indeed, in *T. brucei* RPA foci persist throughout the cell cycle following DNA damage, whilst the foci are only seen in S and G2 phase in mammals (Glover et al. 2019).

Protein phosphorylation is a dynamic process (Gelens et al. 2018) and our results show only a snapshot of the DDR capturing the response at 12 h post DSB induction. Other post translational modifications are also important to both the DDR (Cremona, Sarangi, and Zhao 2012; Lee et al. 2018; Van and Santos 2018) and *VSG* switching, with histone methylation and acetylation important for antigenic variation (Figueiredo, Janzen, and Cross 2008; Glover and Horn 2014; Wang, Kawahara, and Horn 2010), and summoylated proteins enriched at the active BES (Lopez-Farfan et al. 2014). This study is the first DNA damage phosphoproteome in *T. brucei*, and we have identified an abundance of novel proteins involved in the DDR. Validation of candidate phosphorylation sites from this dataset will provide key insights into the protein modifications that govern both DSBR and antigenic variation ins *T. brucei*.

## Data availability

The mass spectrometry proteomics data have been deposited to the ProteomeXchange Consortium via the PRIDE (Perez-Riverol et al. 2022) partner repository with the dataset identifier PXD034455. The the data for this study have been deposited in the European Nucleotide Archive (ENA) at EMBL-EBI under accession number PRJEB52305 (ERP137006).

## Acknowledgements

EJM, MM, MU and LG designed the experiments. EJM, ADH, TC performed the experiments, EJM, QGG and MU performed the statistical analysis. EJM and LG wrote the manuscript, all authors edited the manuscript. We would also like to thank Sebastian Hutchinson for his guidance with the analysis of the VSG-sequencing.

## Supplementary Figure legends

**Supplementary Figure 1.**
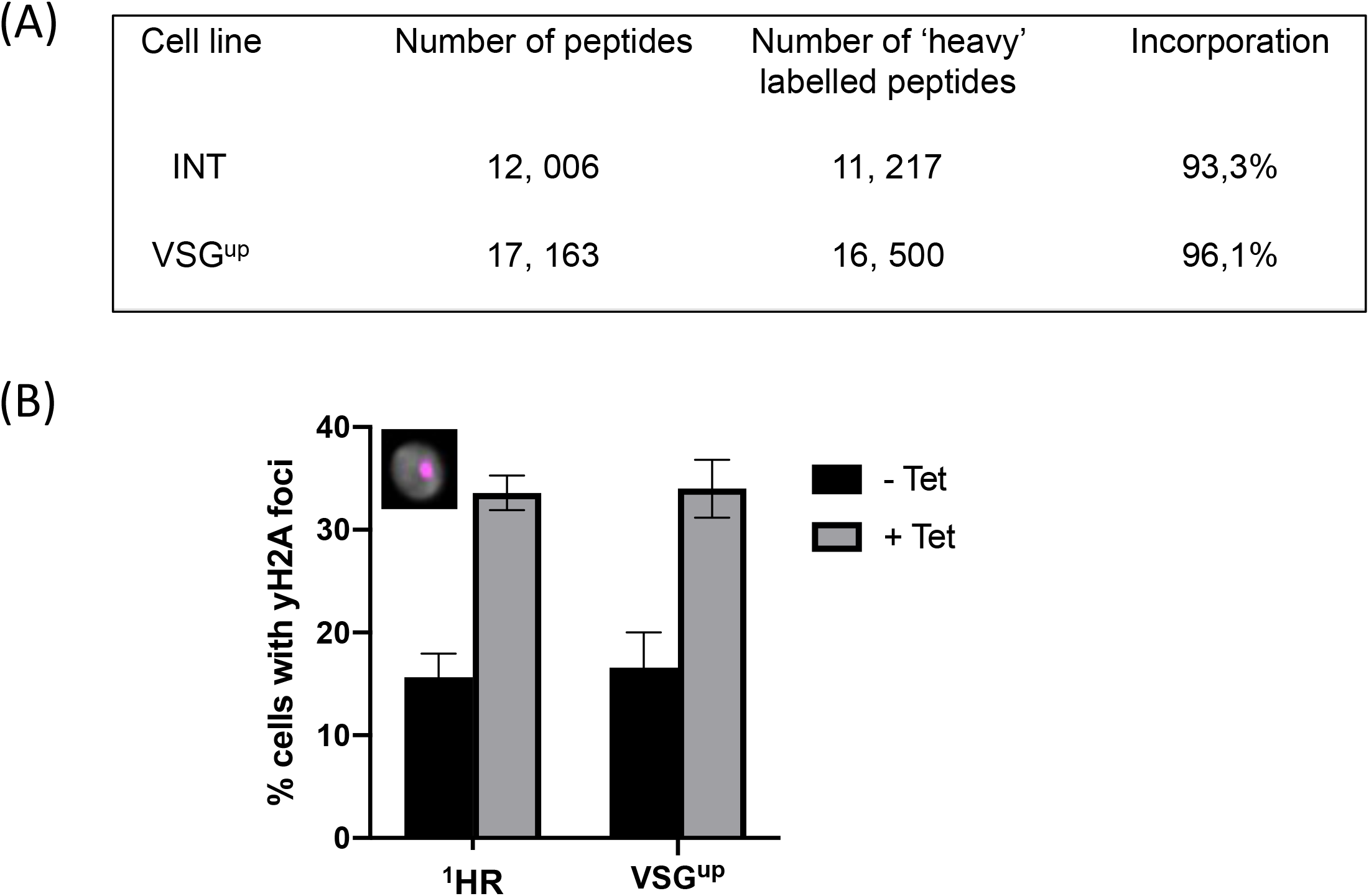
Adaptation of the ^1^HR and VSGup cell lines to growth in SILAC medium. (A) Incorporation of heavy and light labels. (B) γH2A foci formation in cell lines adapted to growth in SILAC HMI-9 medium. ‘-Tet’ indicates uninduced cells, ‘+ Tet’ indicates 12 h DSB induction. n > 100 for each count, and the error bars are the standard deviation between cells grown in ‘heavy’ and ‘light’ SILAC HMI-9. Inset, example of a γH2A positive nuclei, with γH2A shown in magenta and the DAPI nuclear stain in grey. Scale bar, 2 μm.

**Supplementary Figure 2.**
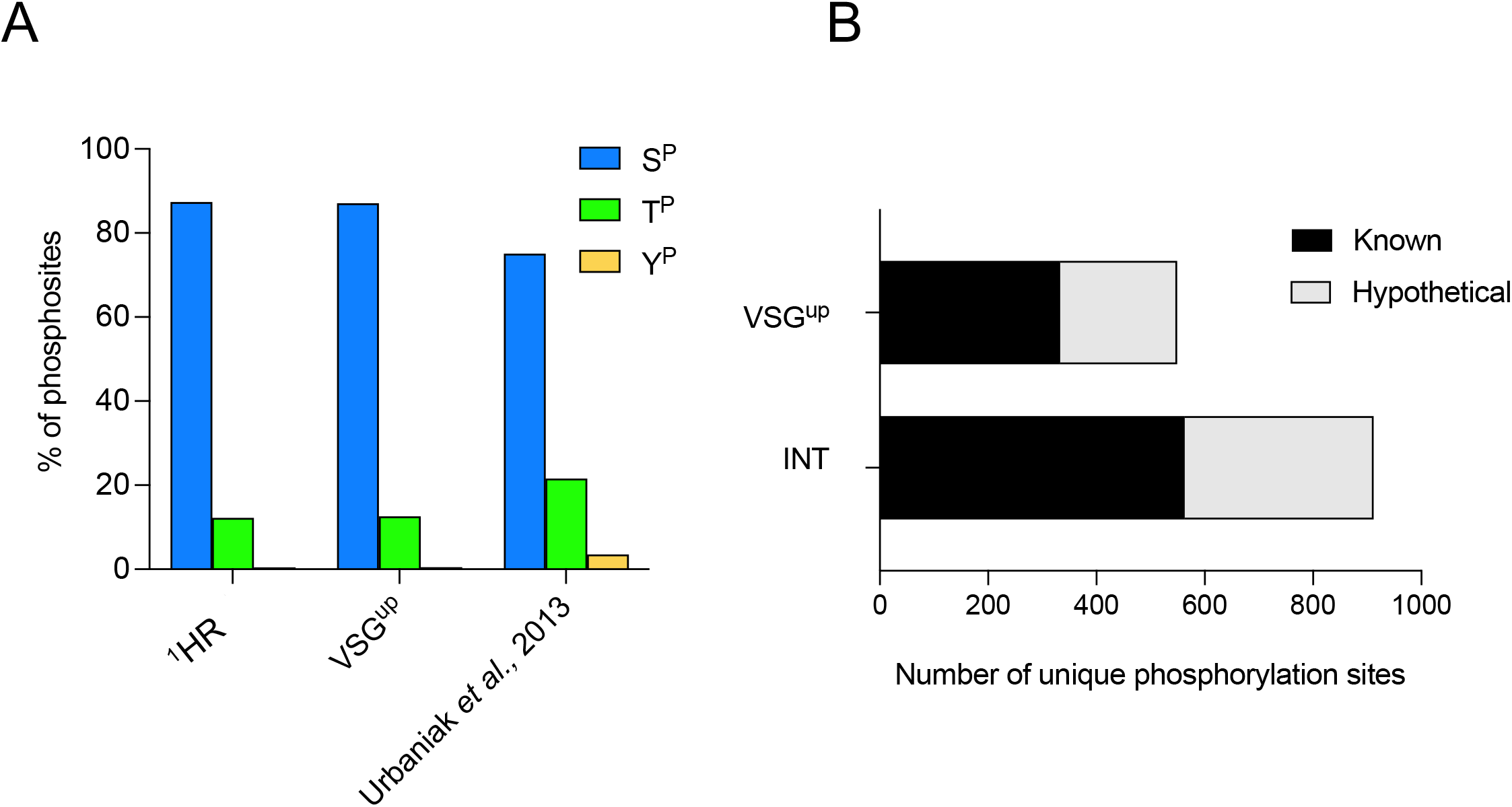
*T. brucei* DNA damage phosphoproteome. (A) Distribution of the total phosphorylation sites identified in the ^1^HR and VSG^up^ DNA damage phosphoproteomes amongst phospho serine (p^S^), thereonine (p^T^) and typrosine (p^Y^). A comparison to the published global phosphoproteome of *T. brucei* is shown (Urbaniak et al. 2013). (B) Number of phosphorylation sites in the ^1^HR and VSG^up^ phosphoproteomes that are located on proteins identified as hypothetical or hypothetical conserved.

**Supplementary Figure 3.**
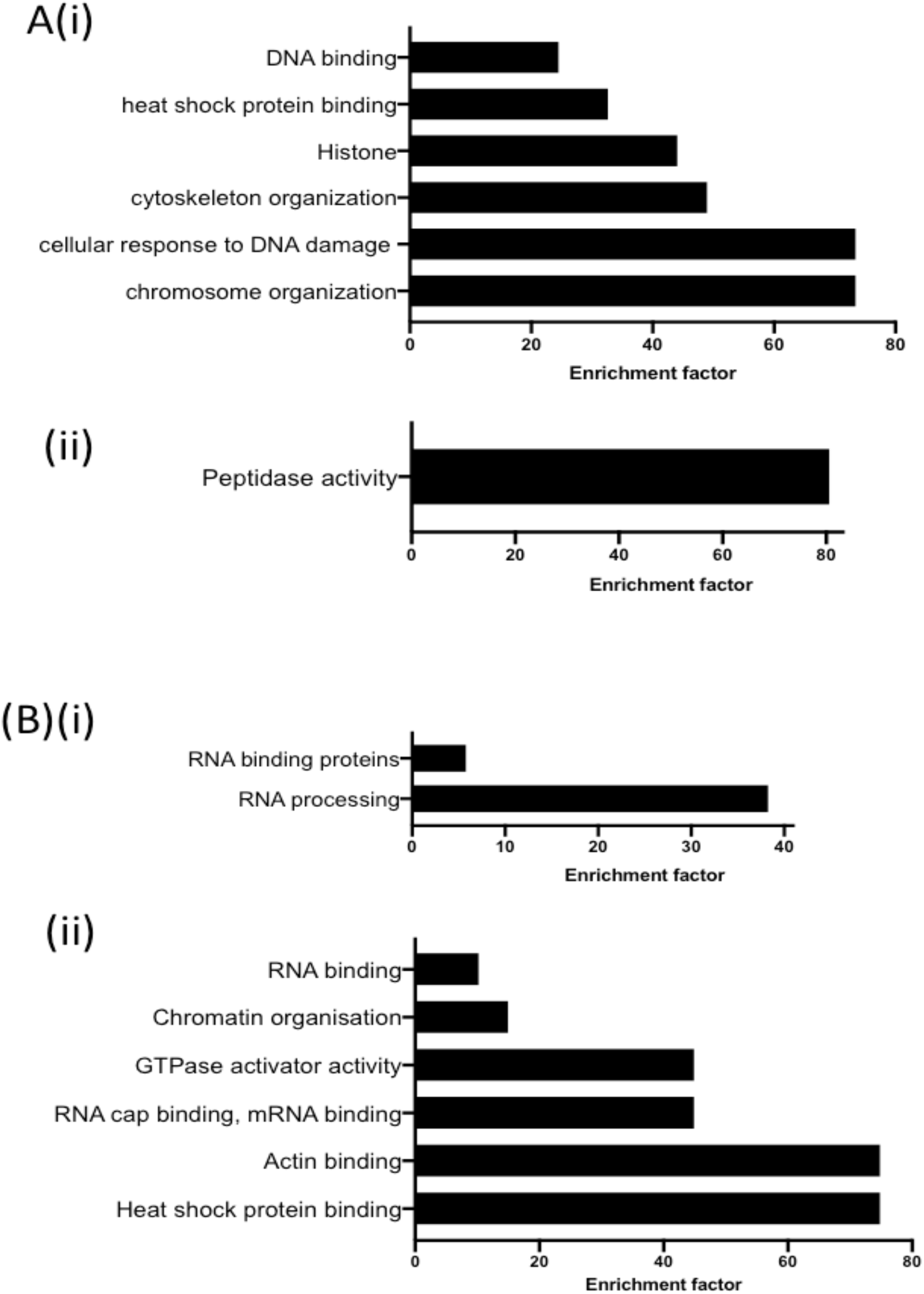
Categorical enrichment of GO terms DNA damage responsive phosphoproteins. (A) GO terms enriched in proteins whose phosphorylation increases following (i) an ^1^HR DSB and (ii) a VSG^up^ DSB. (B) GO terms enriched in proteins whose phosphorylation decreases following (i) an ^1^HR DSB and (ii) a VSG^up^ DSB.

**Supplementary Figure 4.**
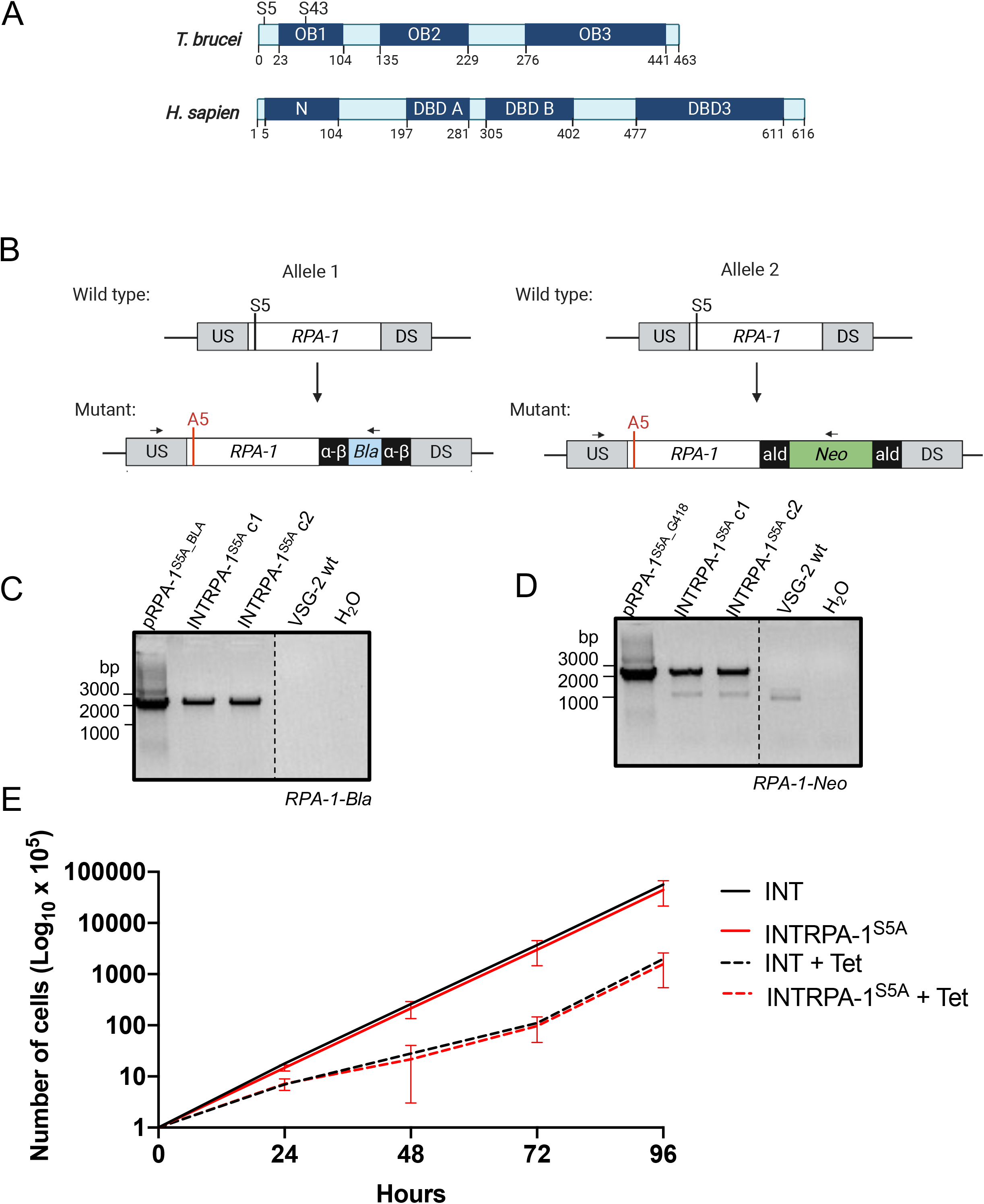
Generation of ^1^HR RPA-1^S5A^ phosphorylation mutants. (A) Schematic showing the major predicted functional domains of *T. brucei* RPA-1 (Tb 927.11.9130) and *H. sapiens* RPA-1 (GI: 4506583) as predicted by InterPro from the amino acid sequences. (B) Schematic of RPA-1 phosphorylation mutant strategy. Top panel shows both wild type *RPA-1* alleles, with the native seine in position 5 highlighted with a black horizontal bar. US and DS refer to the upstream and downstream regions that were targeted for integration of the mutant allele. The US targeting region is 391 bp in length and DS is 498 bp in length for both alleles. Bottom panel shows the mutated *RPA-1* allele with alanine in position 5 shown as a red bar. Allele 1 is replaced with RPA-1S5A fused to a *blasticidin* resistance gene, and Allele 2 is replaced with RPA-1 fused to a *neomycin phosphotransferase* gene. Bla*; blasticidine* resistance gene, a-b; *alpha-beta tubulin* intergenic regions, ald; *aldolase* processing sequences, Neo; *neomycin phosphotransferase resistance* gene Black arrows show regions amplified by PCR to validate integration of the cassette Figure created with BioRender.com. (C) PCR validation of integration of replacement of *RPA-1* allele 1 with the *RPA-1^S5A^ – Bla* in the INT cell line. pRPA-1^S5A^ is the plasmid harbouring the mutant and serves as the positive control. INT*RPA-1*^S5A^ c1 and c2 refer to a pair of biological clones, VSG-2 wt is gDNA extracted from a wild type cell line and serves as a negative control. Expected size of the PCR product is 2246 bp. (D) PCR validation of integration of replacement of *RPA-1* allele 2 with the *RPA-1^S5A^ – Neo* in the INT cell line. Samples as described in (B). Expected size of the PCR product is 2333 bp. (E) Cumulative growth of the INT and INTRPA-1^S5A^ cell lines over 96 h. INT and INTRPA-1^S5A^ cell lines are shown in black and red lines, respectively, and dashed lines indicate growth under DSB inducing conditions. Error bars represent standard deviation between a single induction in a pair of biological clones.

**Supplementary Figure 5.**
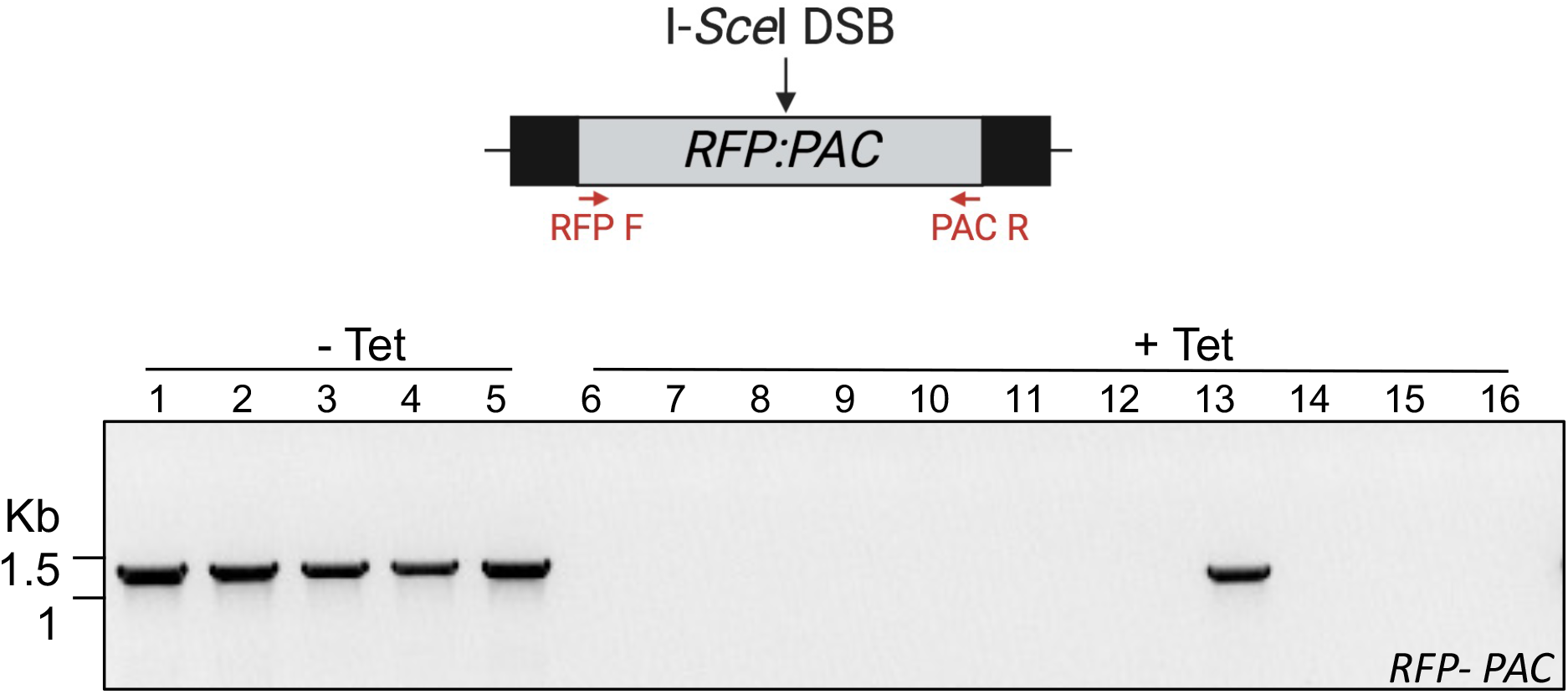
Analysis of repaired ^1^HR RPA-1^S5A^ clones. Above, schematic showing binding site of primers used to assess INT DSB repair. Below, PCR analysis of 5 uninduced subclones (‘-Tet’) and 11 induced subclones (‘+Tet). Data for 9 induced subclones not shown. Expected size of PCR product in uninduced clones is 1.314 Kb.

**Supplementary Figure 6.**
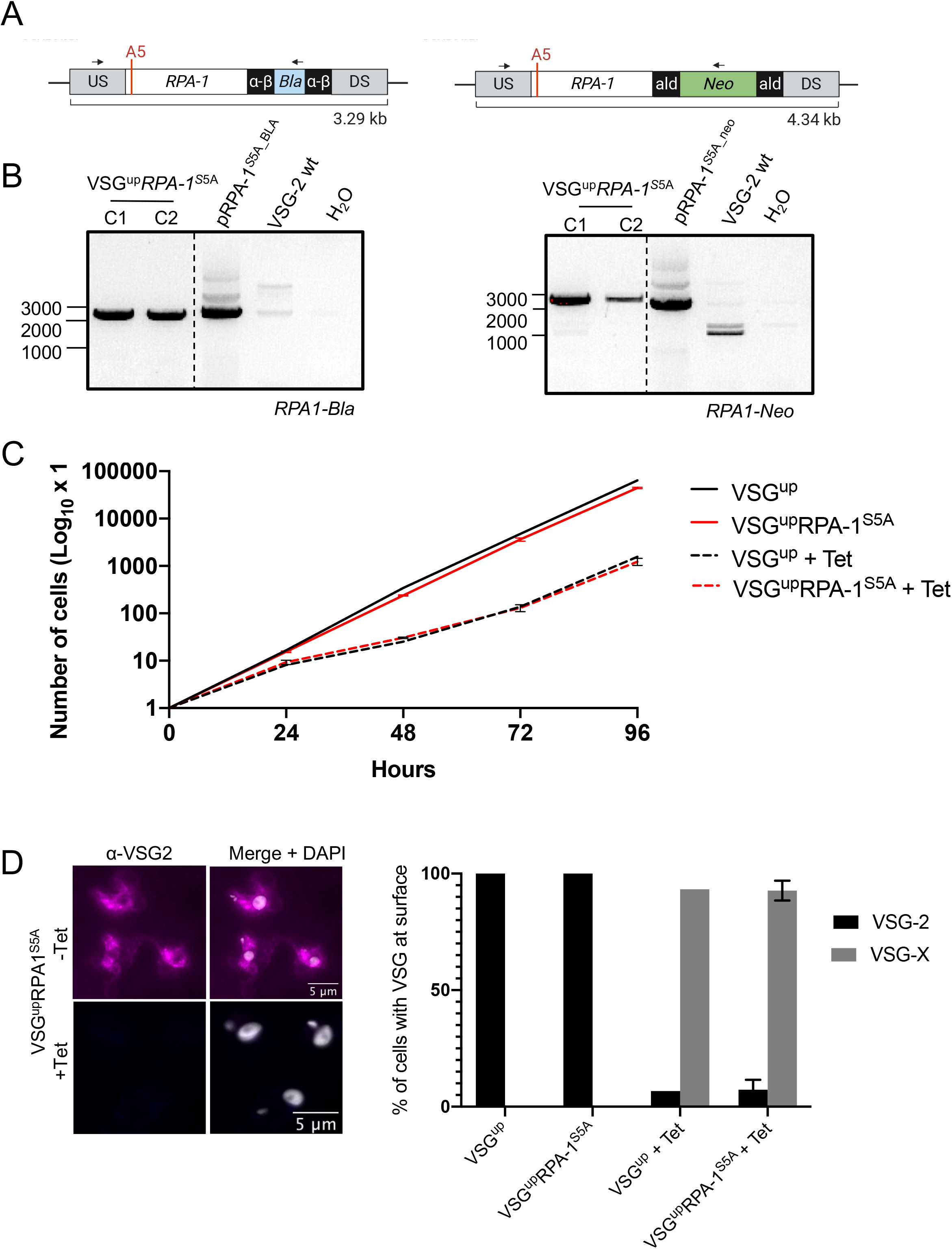
Generation of VSG^up^RPA-1^S5A^ phosphorylation mutants. (A) PCR to confirm replacement of one RPA-1 allele with RPA-1^S5A^ fused to a *blasticidin* resistance cassette in the VSG^up^ cell line. Primer binding sites used for the PCR shown are shown as black arrows. C1 and C2 refer to a pair of VSG^up^RPA-1^S5A^ phosphorylation mutants, pRPA-1S5A is the plasmid and used a a positive control, VSG-2 WT is an unmodified cell line used as negative control. Expected size of the PCR product is 2246 bp. (B) PCR to confirm replacement of the second *RPA-1* allele with *RPA-1*S5A fused to a Neomycin resistance gene in the VSG^up^ cell line. pRPA-1^S5A^ is the plasmid harbouring the mutant and serves as the positive control. Primer binding sites used for the PCR shown are shown as black arrows. Expected size of the PCR product is 2333 bp. (C) Cumulative growth of the VSG^up^ and VSG^up^RPA-1^S5A^ cell lines over 96 h. VSG^up^ and VSG^up^RPA-1^S5A^ cell lines are shown in black and red lines, respectively and dashed lines indicate growth under DSB inducing conditions. Error bars represent standard deviation between a single induction in a pair of biological clones. (D) Immunofluorescence analysis of surface VSGs in the VSG^up^ and VSG^up^*RPA-1*^S5A^ cell lines, in uninduced and 7 days DSB induced samples (+ Tet). VSG-2 is the surface VSG in the VSG^up^ cell line prior to DSB induction. VSG-X refers to all VSGs except VSG-2 and represents the proportion of cells that no longer have VSG-2 at the surface and have therefore undergone VSG switching. Error bars for VSG^up^*RPA-1*^S5A^ cell line represent standard deviation between a pair of biological clones. n>100 for each sample. Panel on the right shows VSG^up^*RPA-1*^S5A^ cells before DSB induction (-Tet) and 7 days post DSB induction (+ Tet), stained using antibodies specific to VSG-2.

**Supplementary Figure 7.**
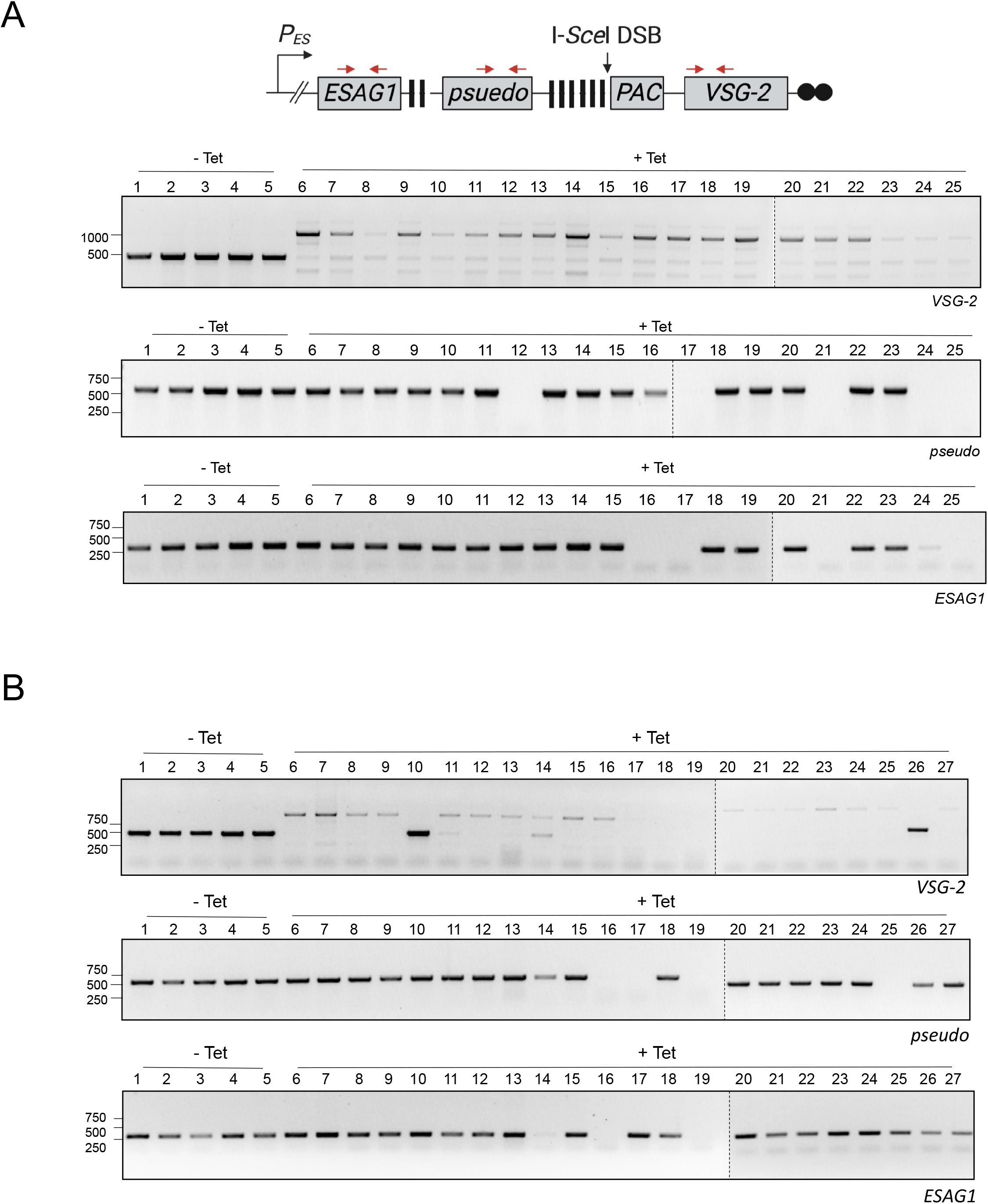
Analysis of repaired VSG^up^RPA-1^S5A^ clones. (A) VSG^up^ subclone analysis of *VSG-2, pseudo* and *ESAG1* which have an expected PCR product size of 527bp, 550bp and 380bp, respectively. Lanes 1-5 are uninduced subclones (-Tet) and lanes 6 – 25 are induced (+ Tet). (B) VSG^up^*RPA-1*^S5A^ subclone analysis of VSG*-2, pseudo* and *ESAG1.* Lanes 6 -27 are induced subclones.

